# Downregulation of insulin-like growth factor binding protein 5 is involved in intervertebral disc degeneration via the ERK signaling pathway

**DOI:** 10.1101/425272

## Introduction

Intervertebral disc degeneration (IDD) is a disease of discs connecting adjoining vertebrae in which structural damage causes the degeneration of the disc and the surrounding area. This disease represents a common cause of back pain, and its treatment is costly and relatively ineffective (Kepler et al., 2013), with patients who do not achieve improvement with conservative management possibly requiring surgery (Lurie et al., 2014; Postacchini, 1996). It has been reported that the central features of IDD are the reduction in the nucleus pulposus (NP) cell population and the loss of extracellular matrix molecules (ECM), which eventually leads to major changes in the architecture and properties of the disc (Tang et al., 2016).

The survival of NP cells in the avascular nondegenerated disc is likely attributed to cell-secreted growth factors that act on the same cells in an autocrine manner. In addition, the expression of several growth factors and their respective receptors has been reported in areas of degeneration in human and animal intervertebral discs (Leckie et al., 2012). Thus, a comprehensive understanding of growth factors is essential to maximize opportunities to develop therapeutic interventions that retard or reverse IDD.

There are classical mitogens such as platelet-derived growth factor (PDGF), basic fibroblast growth factor (bFGF), and insulin-like growth factor I (IGF-I). IGF-I and the members of the insulin family of proteins, including insulin-like growth factor binding proteins (IGFBPs), have been reported to concomitantly influence both metabolic and proliferative processes. IGFBP5 is one of the conserved IGFBPs that can translocate into the nucleus, as determined by the presence of a nuclear localization sequence (Yamaguchi et al., 2011). IGFBP5 has been shown to enhance cell growth and the remodeling and repair of bone. Daily subcutaneous injections of recombinant IGFBP5 protein stimulates osteoblast activity and bone accretion in ovariectomized mice (Andress, 2001). Moreover, a previous study demonstrated that IGFBP5 plays a crucial role in the carcinogenesis and progression of several types of cancer, such as oral and breast cancer, and its expression is usually dysregulated (Chang et al., 2010; Mita et al., 2007; Wang et al., 2015). However, the effect of IGFBP5 on regulating proliferative and apoptotic activities in NP cells is not well defined, even though it is expressed in NP cells.

Extracellular signal-regulated kinase (ERK) is an extremely conserved signaling pathway that is crucially involved in cell proliferation and survival and diverse cellular processes (Namkoong et al., 2011; Risbud and Shapiro, 2014). A previous study also demonstrated that exogenous and autocrine growth factors stimulate human intervertebral disc cell proliferation via the ERK and Akt pathways (Pratsinis et al., 2012). However, the correlation between IGFBP5, the ERK signaling pathway, and IDD have not been elucidated. Therefore, we aimed to determine the role of IGFBP5 in NP cell proliferation and apoptosis, and the involvement of the ERK signaling pathway in this process.

## Results

### Identification of mRNAs differentially expressed in degenerative NP tissues

The mRNA expression profiles were detected in 10 IDD tissues and 10 normal samples, and Solexa sequencing was performed to show distinguishable mRNA expression patterns among the samples. From the mRNA expression profiling data, 89 mRNAs were found to be differentially expressed in IDD tissues, of which 41 mRNAs were upregulated and 48 mRNAs were downregulated. These differentially expressed serum miRNAs were chosen for further study only when they met the following criteria: (1) having at least 20 copies of mRNA expression; (2) mean fold change >2.0 or <0.5; and (3) p values <0.05. Based on these criteria, 23 mRNAs, of which 14 mRNAs were upregulated and 9 mRNAs were downregulated in patients compared with controls, were chosen for further validation (Fig. 1A).

**Fig. 1.**
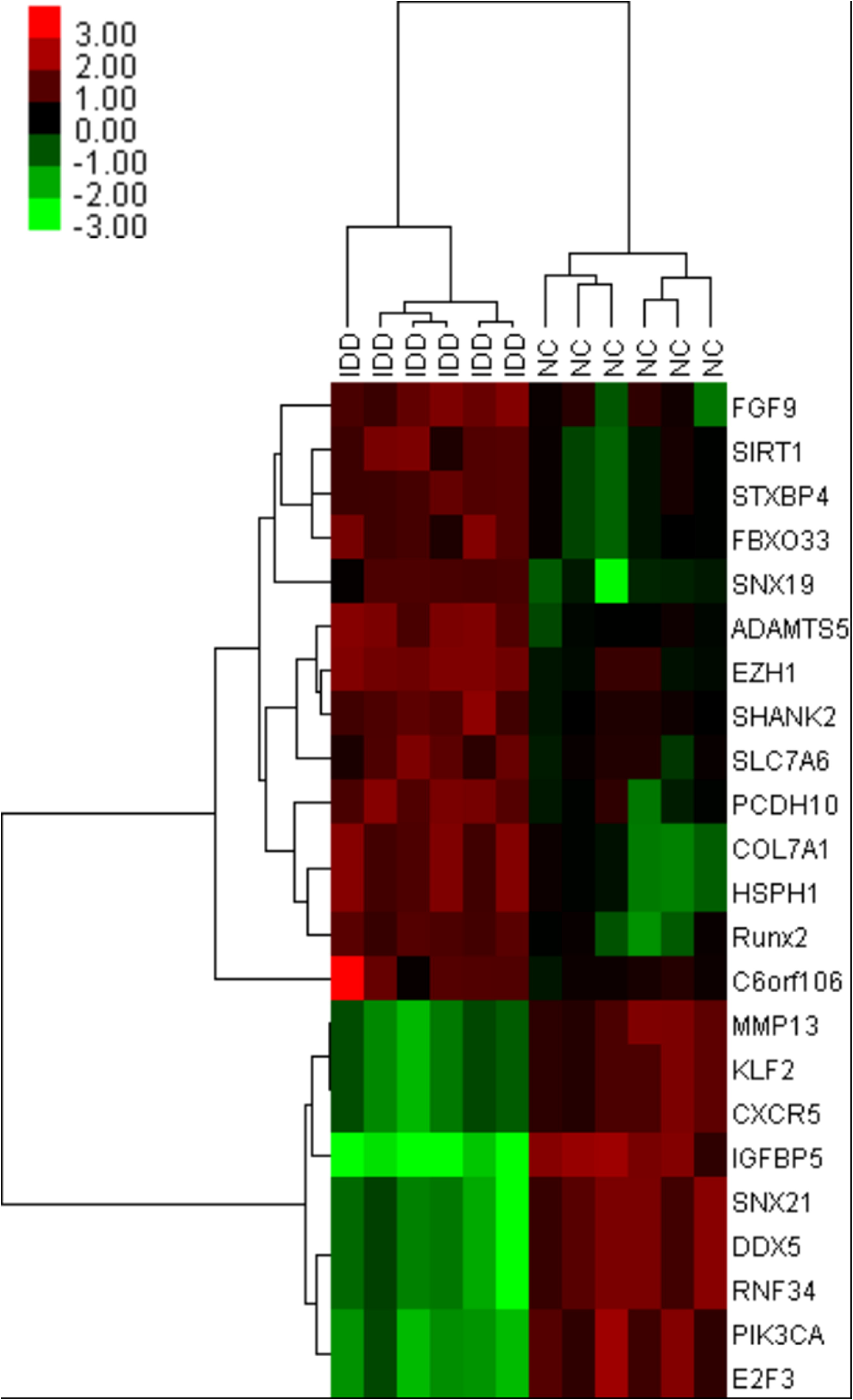
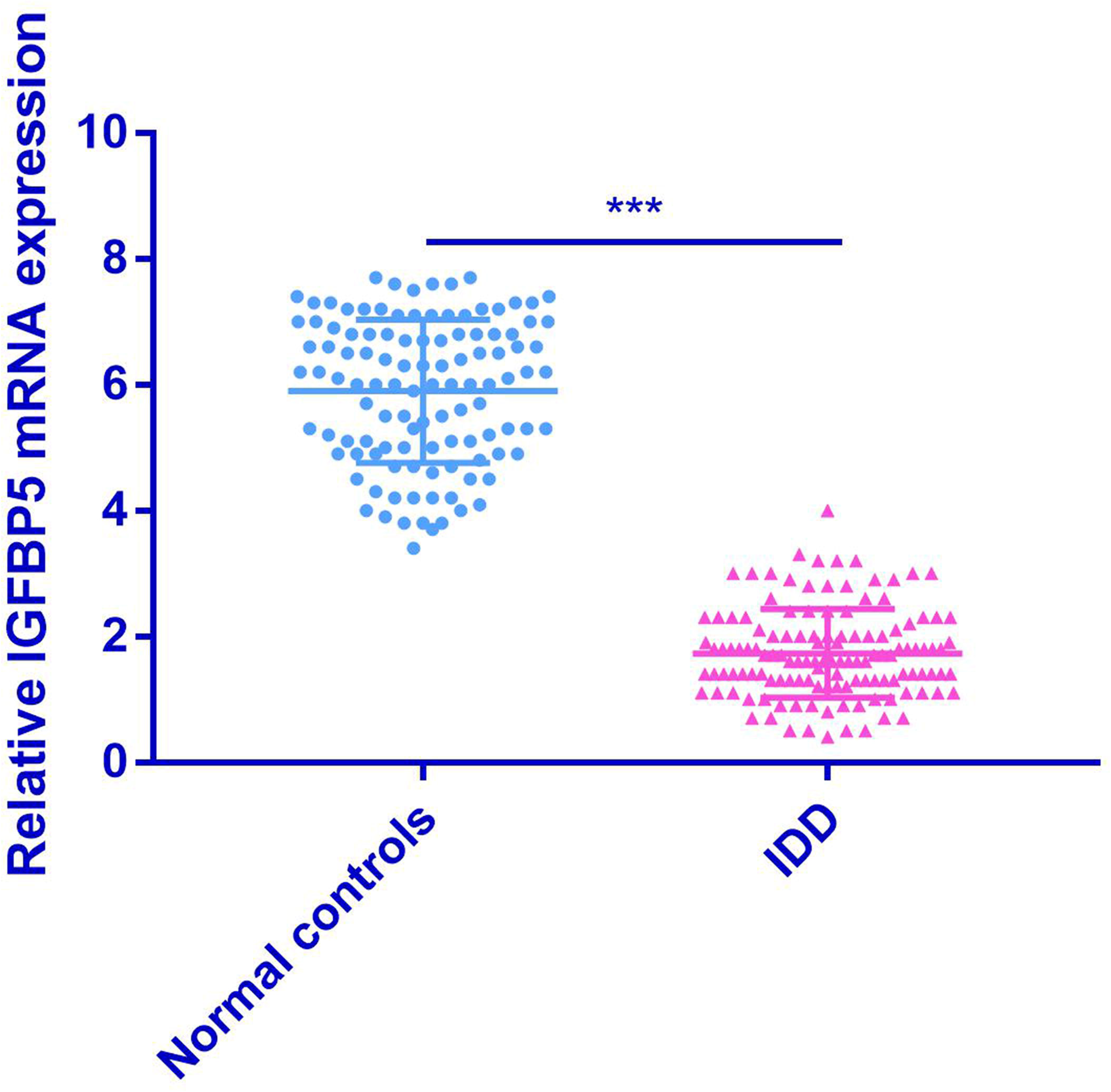
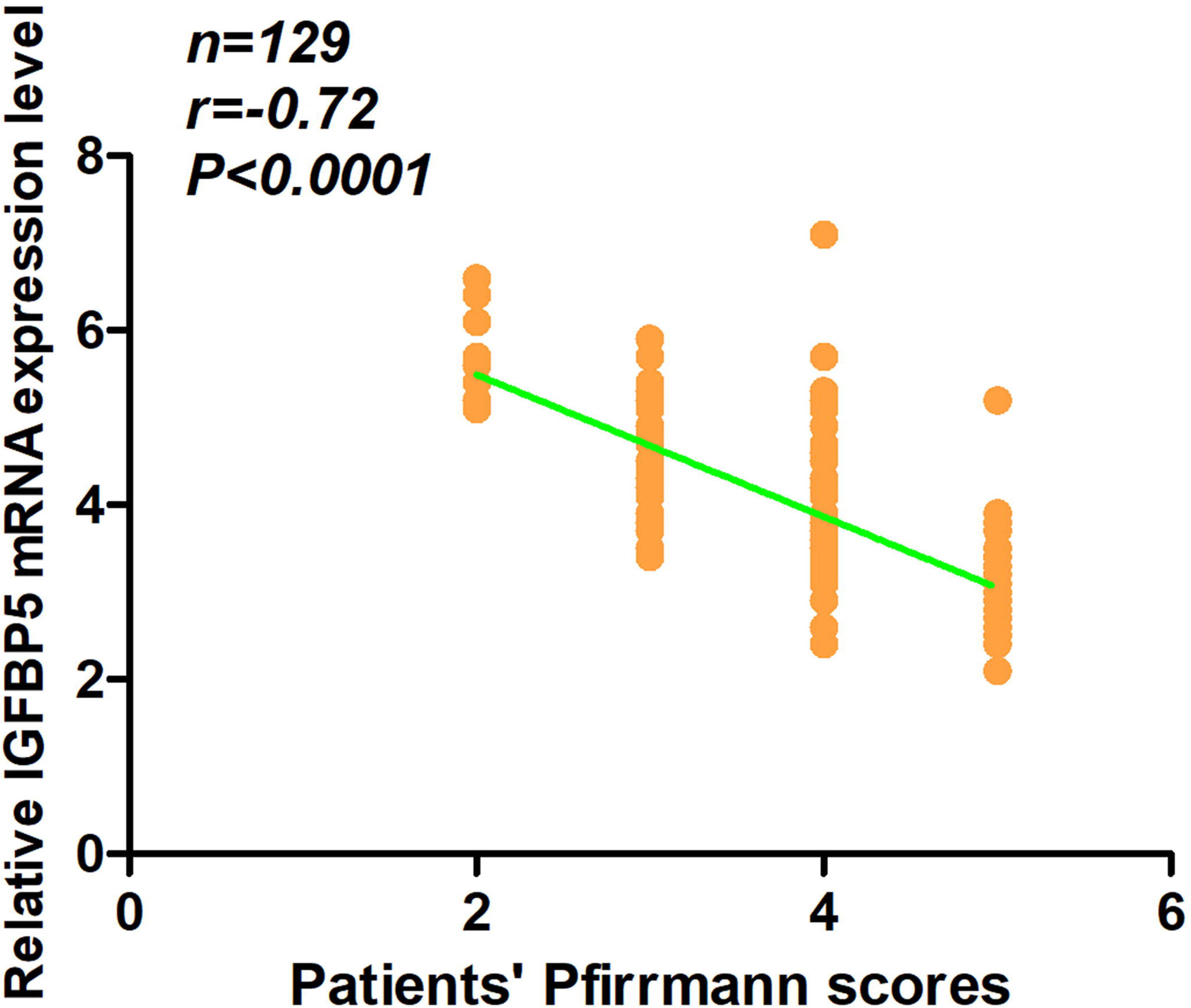
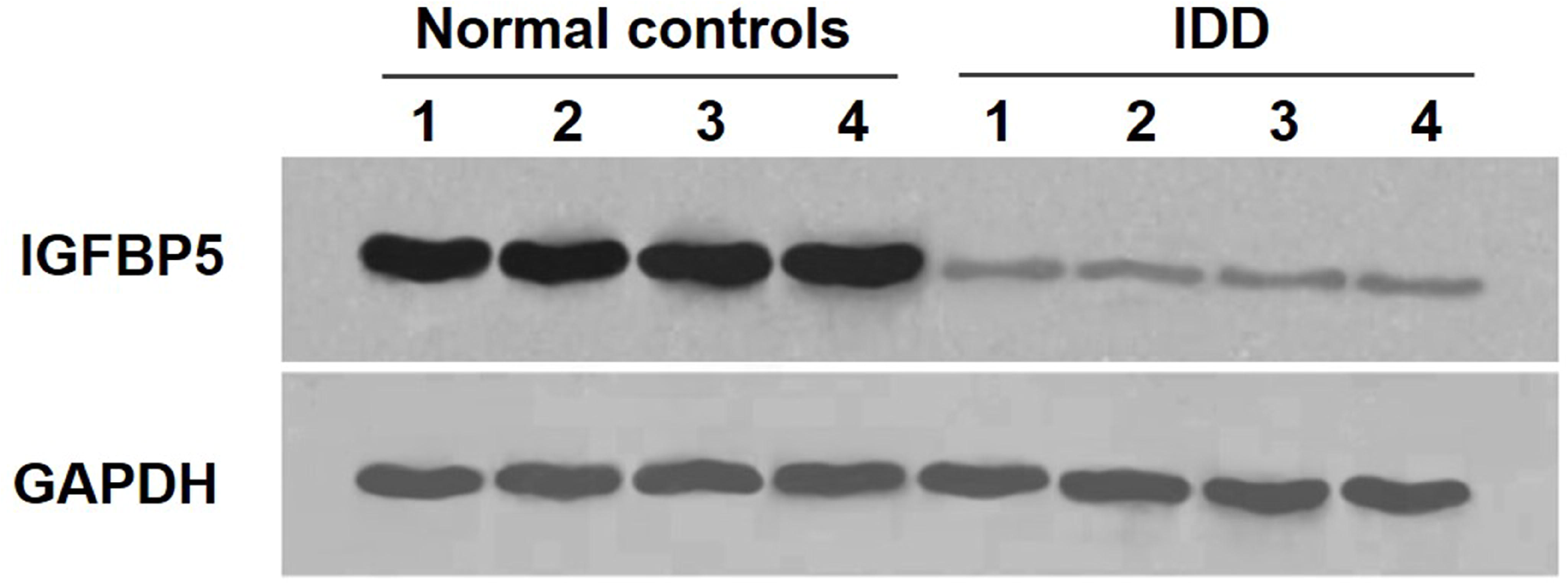
Identification of mRNAs differentially expressed in degenerative NP tissues. (A) A heat map was generated by unsupervised clustering analyses with 23 significantly dysregulated mRNAs in patients with IDD. Hierarchical clustering was performed with average linkage and uncentered correlation. The mRNA expression profile effectively segregated patients with IDD from NCs. (B) The expression level of IGFBP5 mRNA in NP cells was measured in 129 patients and 112 NCs (in the training and validation sets) using RT-qPCR assay (***p < 0.001). (C) The IGFBP5 mRNA expression level was inversely correlated with the Pfirrmann scores (r=-0.72, p < 0.0001). (D) The expression of IGFBP5 in degenerative NP tissues was significantly lower than in normal NP samples, as assessed by Western blots analysis (p < 0.05). IDD=intervertebral disc degeneration; NCs=normal controls; NP=nucleus pulposus.

RT-qPCR assay was used to confirm the expression of the candidate mRNAs. In the training set, mRNAs were measured in a separate set of samples from 10 patients and 10 controls of the previous step. Only mRNAs with a mean fold change >2.0 or <0.5 and a p value <0.01 were selected for further analysis. Using the abovementioned criteria, SIRT1, PCDH10, IGFBP5, and PIK3CA were observed to be significantly different in patients compared with controls (Table 1). In the validation set, the concentration of SIRT1, PCDH10, IGFBP5, and PIK3CA were measured by RT-PCR in a larger cohort comprised of 129 patients and 112 controls. The IGFBP5 expression pattern in the validation set was consistent with those in the training set. Compared with the controls, the level of IGFBP5 mRNA was significantly lower in patients (Table 1). Thus, we focused on IGFBP5 mRNA for further study.

**Table 1.** Differentially expressed mRNAs in IDD compared with normal controls in both the training set and the validation set.

| mRNA | Training set |  | Validation set |  |
| --- | --- | --- | --- | --- |
|  | Fold change | P Value | Fold change | P Value |
| **Upregulated** |  |  |  |  |
| FGF9 | 4.1 | 0.23 | - | - |
| **SIRT1** | **7.4** | **0.006\*\*** | **7** | **0.15** |
| STXBP4 | 4.1 | 0.32 | - | - |
| FBXO33 | 8.9 | 0.34 | - | - |
| SNX19 | 7.5 | 0.09 | - | - |
| ADAMTS5 | 6.1 | 0.34 | - | - |
| EZH1 | 7.3 | 0.45 | - | - |
| SHANK2 | 6.2 | 0.38 | - | - |
| SLC7A6 | 7.6 | 0.27 | - | - |
| **PCDH10** | **6.6** | **0.004\*\*** | **6.2** | **0.32** |
| COL7A1 | 8.2 | 0.78 | - | - |
| HSPH1 | 7.1 | 0.29 | - | - |
| Rux2 | 4.7 | 0.13 | - | - |
| C6orf106 | 6.2 | 0.07 | - | - |
| **Downregulated** |  |  |  |  |
| MMP13 | 0.19 | 0.14 | - | - |
| KLF2 | 0.18 | 0.26 | - | - |
| CXCR5 | 0.31 | 0.87 | - | - |
| **IGFBP5** | **0.02** | **0.002\*\*** | **0.031** | **0.005\*\*** |
| SNX21 | 0.11 | 0.43 | - | - |
| DDX5 | 0.37 | 0.35 | - | - |
| RNF34 | 0.21 | 0.56 | - | - |
| **PIK3CA** | **0.18** | **0.006\*\*** | **0.11** | **0.32** |
| E2F3 | 0.12 | 0.43 | - | - |

\*\* P < 0.01.

### Downregulation of IGFBP5 expression is associated with IDD

To determine the expression level of IGFBP5 mRNA in degenerative NP tissues, we measured the level of IGFBP5 mRNA in 129 patients and 112 controls. The results demonstrated that the level of IGFBP5 mRNA was downregulated in degenerative NP tissues when compared with controls (Fig. 1B). In addition, the expression of IGFBP5 mRNA was negatively correlated with the disc degeneration grade (r=-0.72, p < 0.0001) (Fig. 1C). In addition, the expression of IGFBP5 in degenerative NP tissues was significantly lower than in normal NP samples according to the Western blots analysis (p < 0.05, Fig. 1D).

### IGFBP5 promotes NP cell proliferation and inhibits apoptosis in vitro

At 10 days following IGFBP5-overexpressing transfection, cell proliferation was significantly higher compared to IGFBP5-shRNA (p < 0.01), as evidenced by the absorbance values at 570 nm (Fig. 2A). The PI staining results revealed that compared with those in the NC group, the percentages of cells in the G1 phase in the IGFBP5-overexpressing and PD98059-treated samples was significantly decreased, while those in the S phase were significantly increased (all P < 0.05, Fig. 2E and 2F). In contrast, IGFBP5 silencing led to cell growth arrest at the G1 phase and the number of cells in the S phase decreased, indicating the suppression of NP cell proliferation (all Ps < 0.05, Fig. 2D). With respect to NP cell apoptosis, the Annexin-V apoptosis assay demonstrated that IGFBP5-overexpressing transfection significantly decreased the number of apoptotic NP cells compared with IGFBP5-shRNA transfection (p < 0.05) (Fig. 2J and 2K). This effect was further confirmed by the colony formation assay (Fig. 2N).

**Fig. 2.**
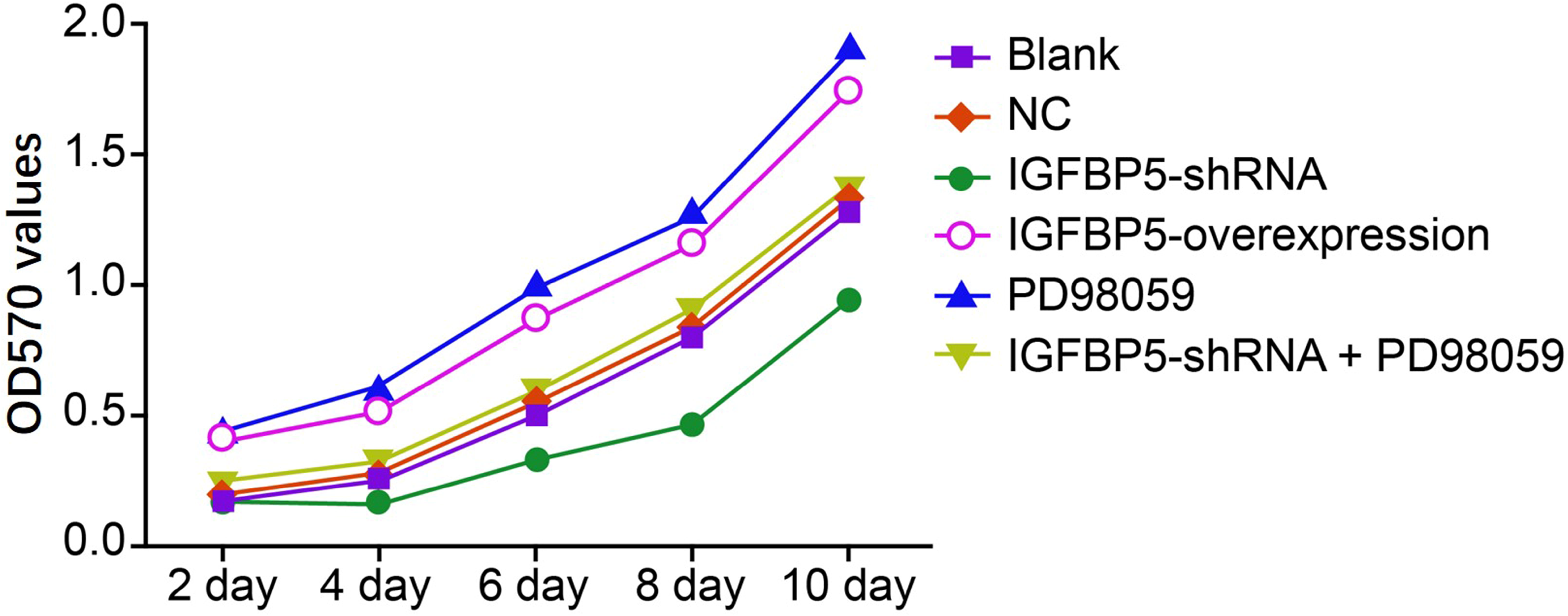
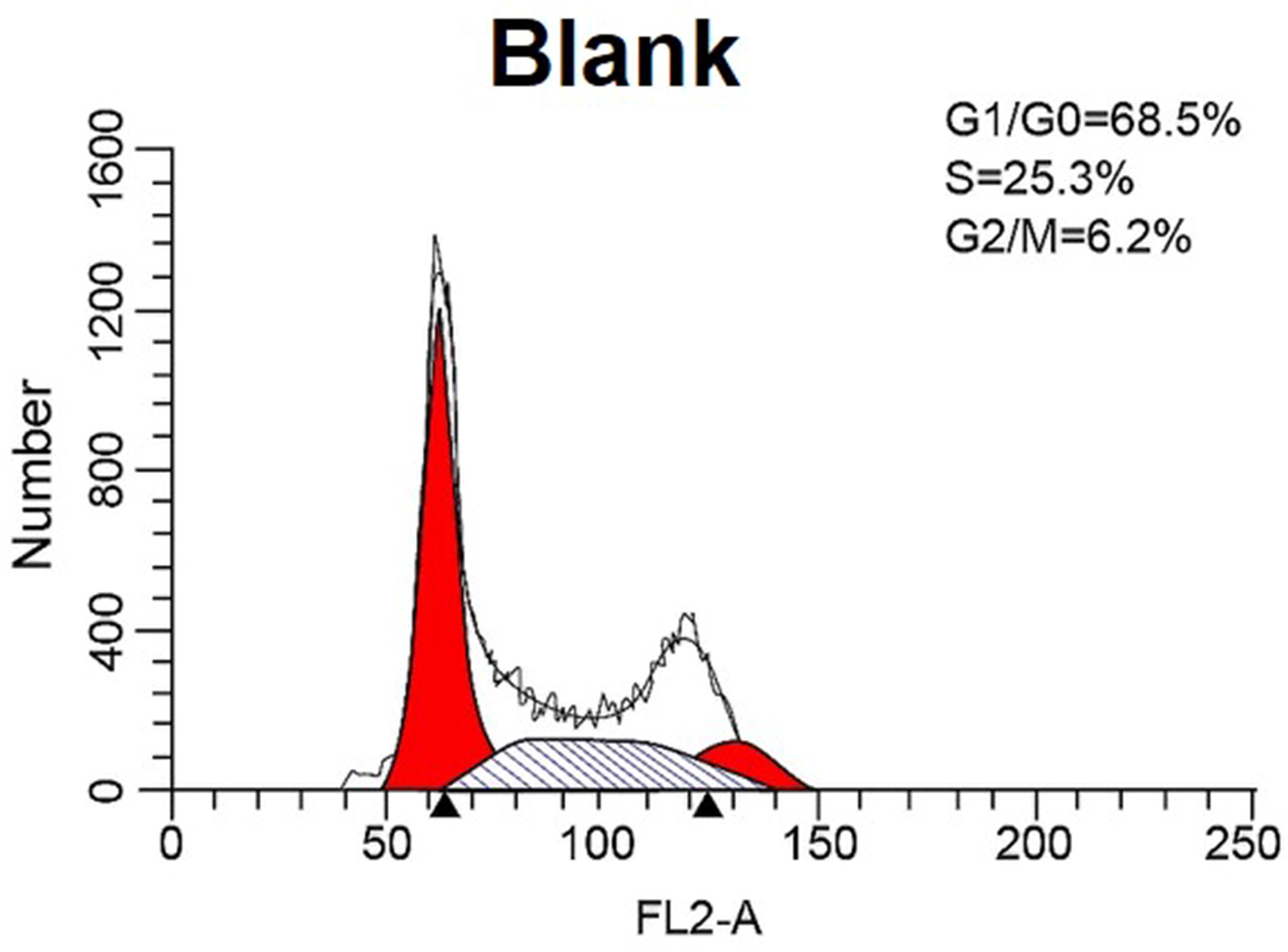
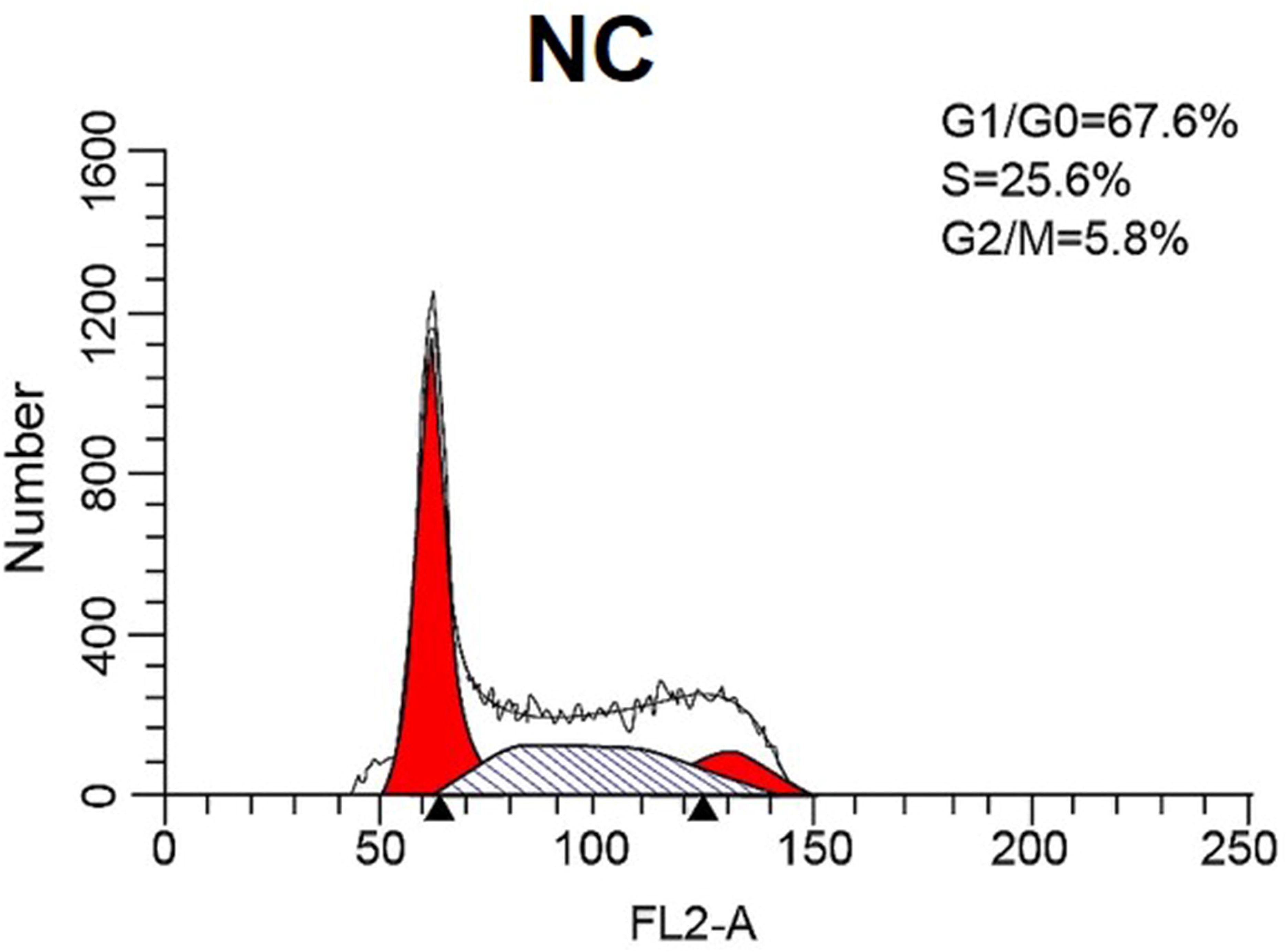
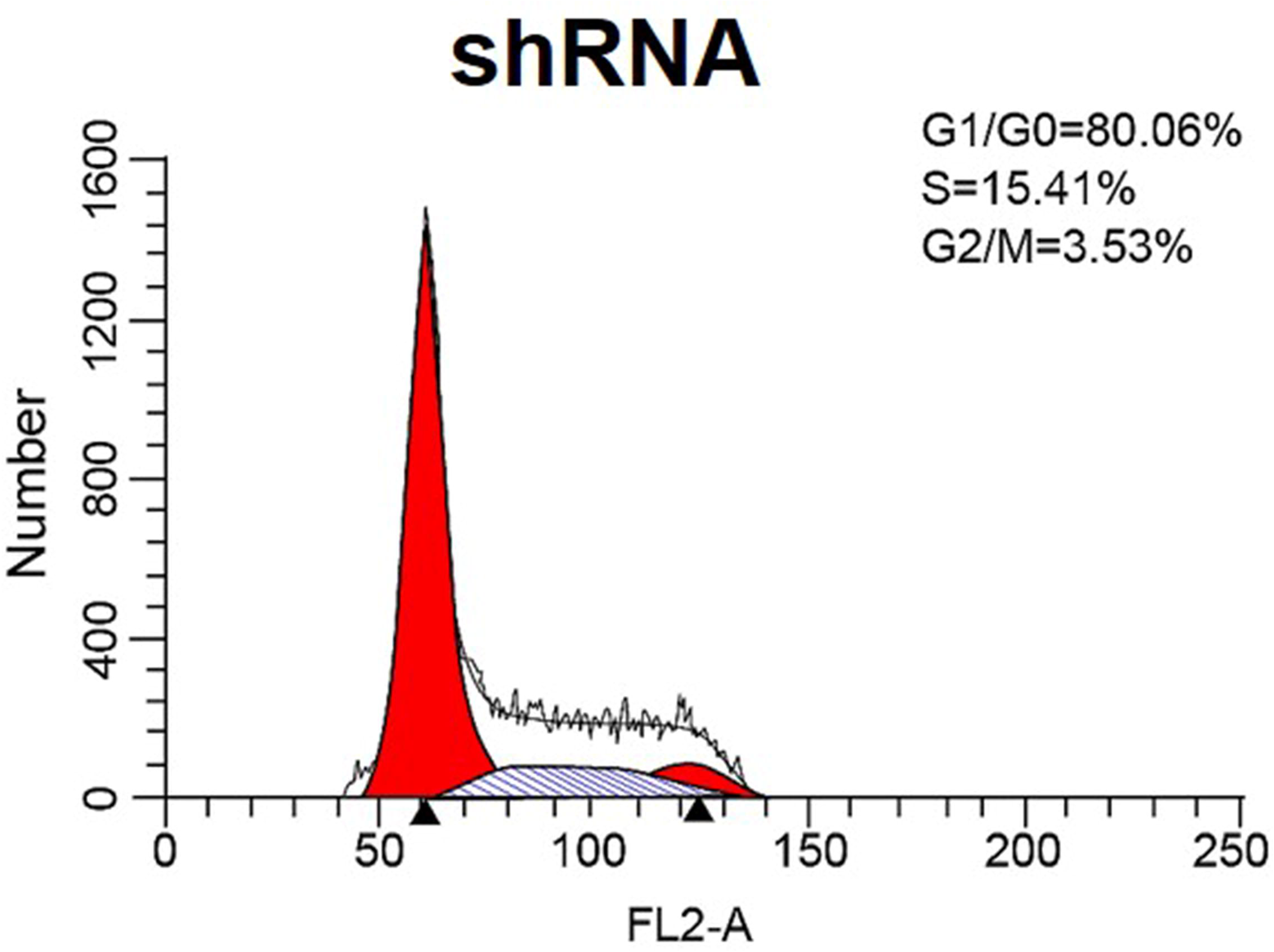
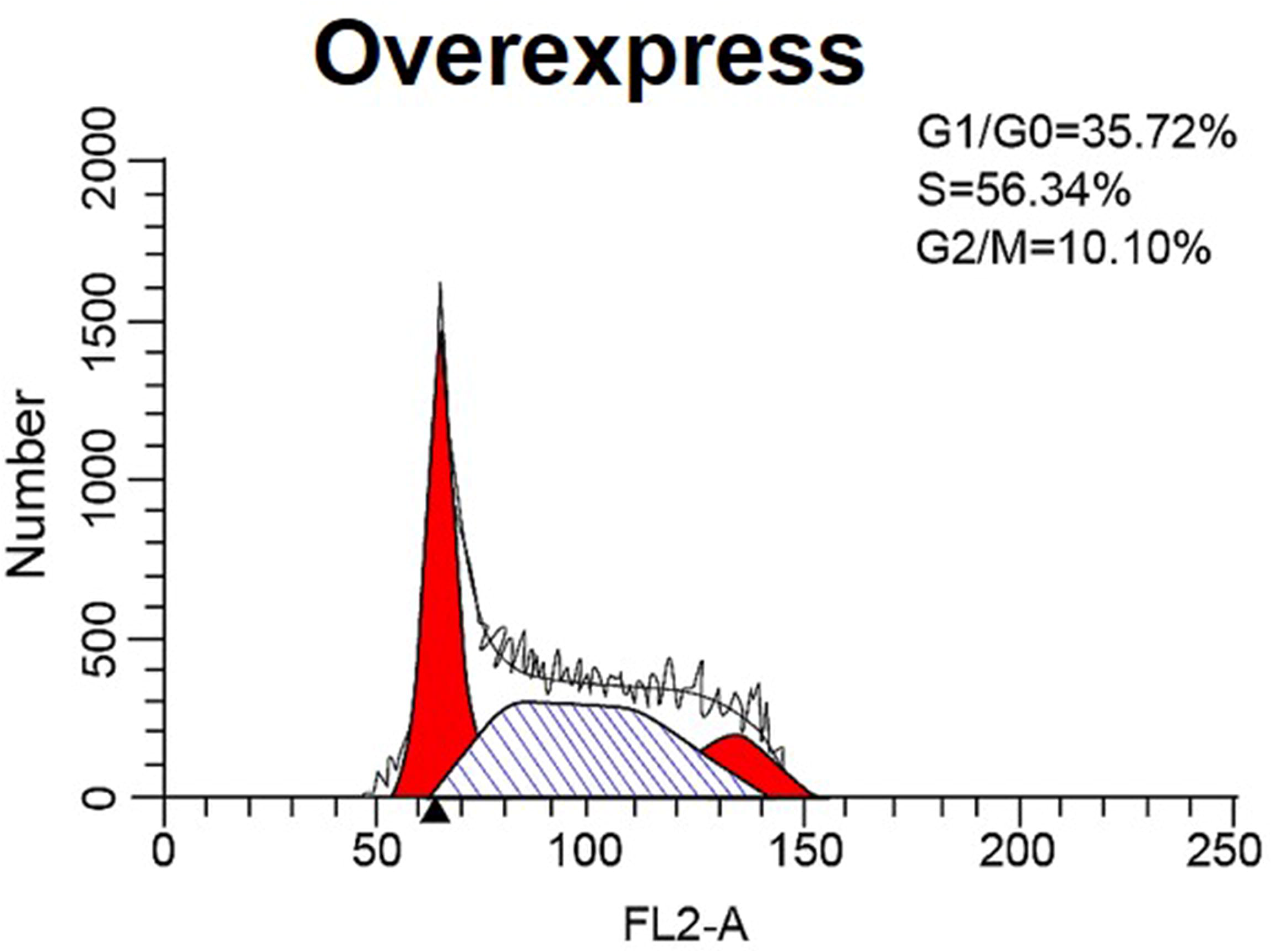
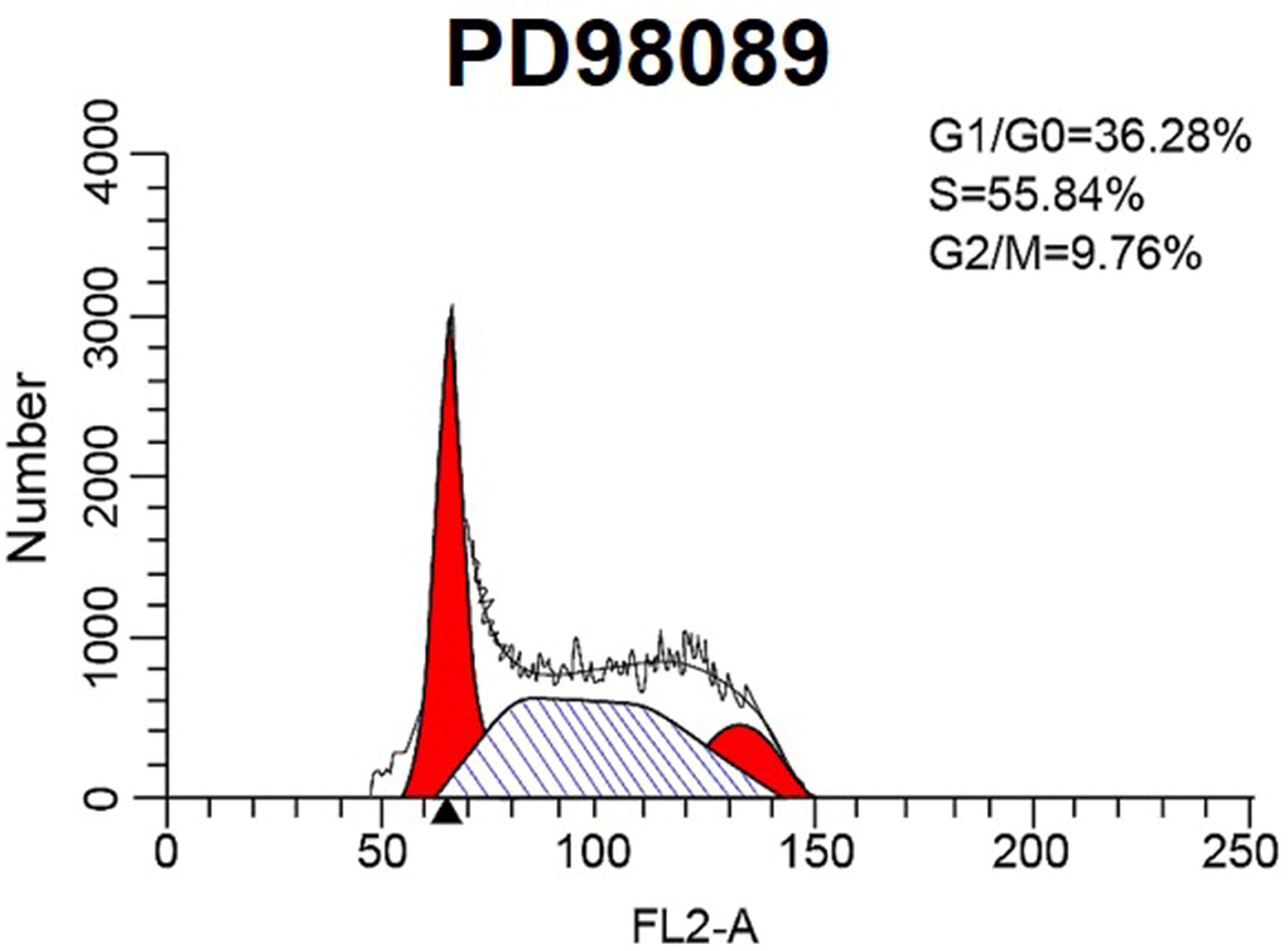
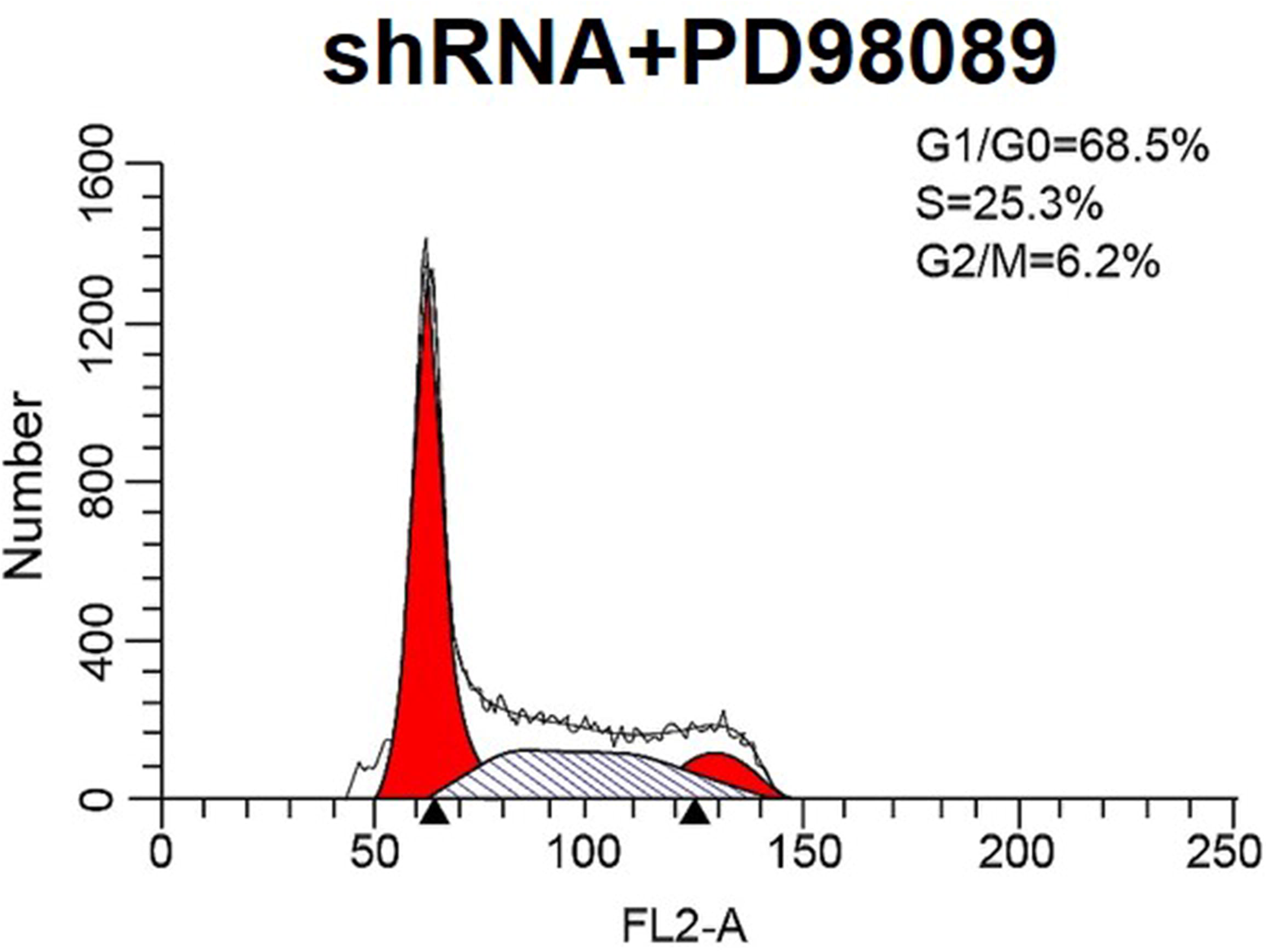
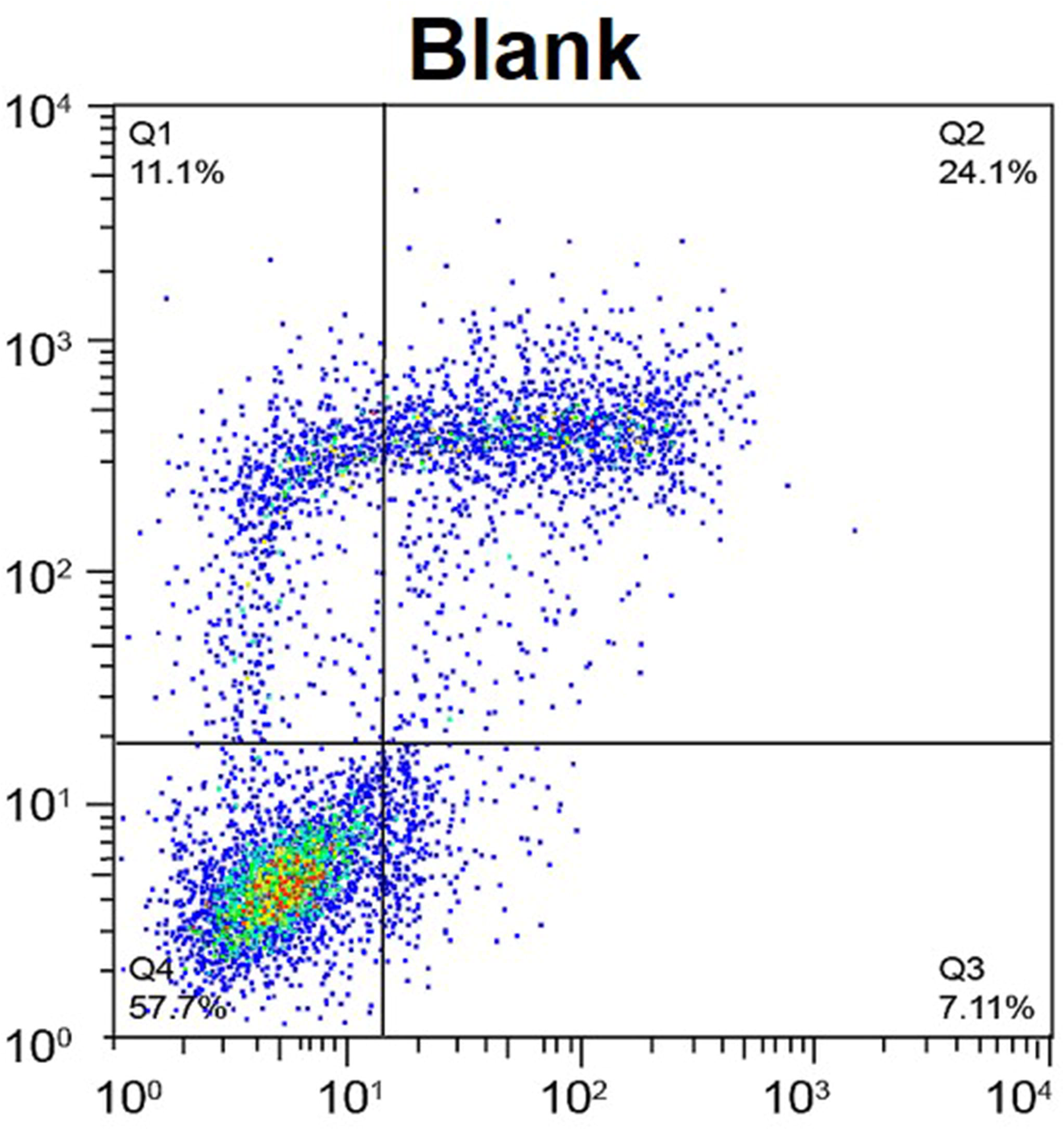
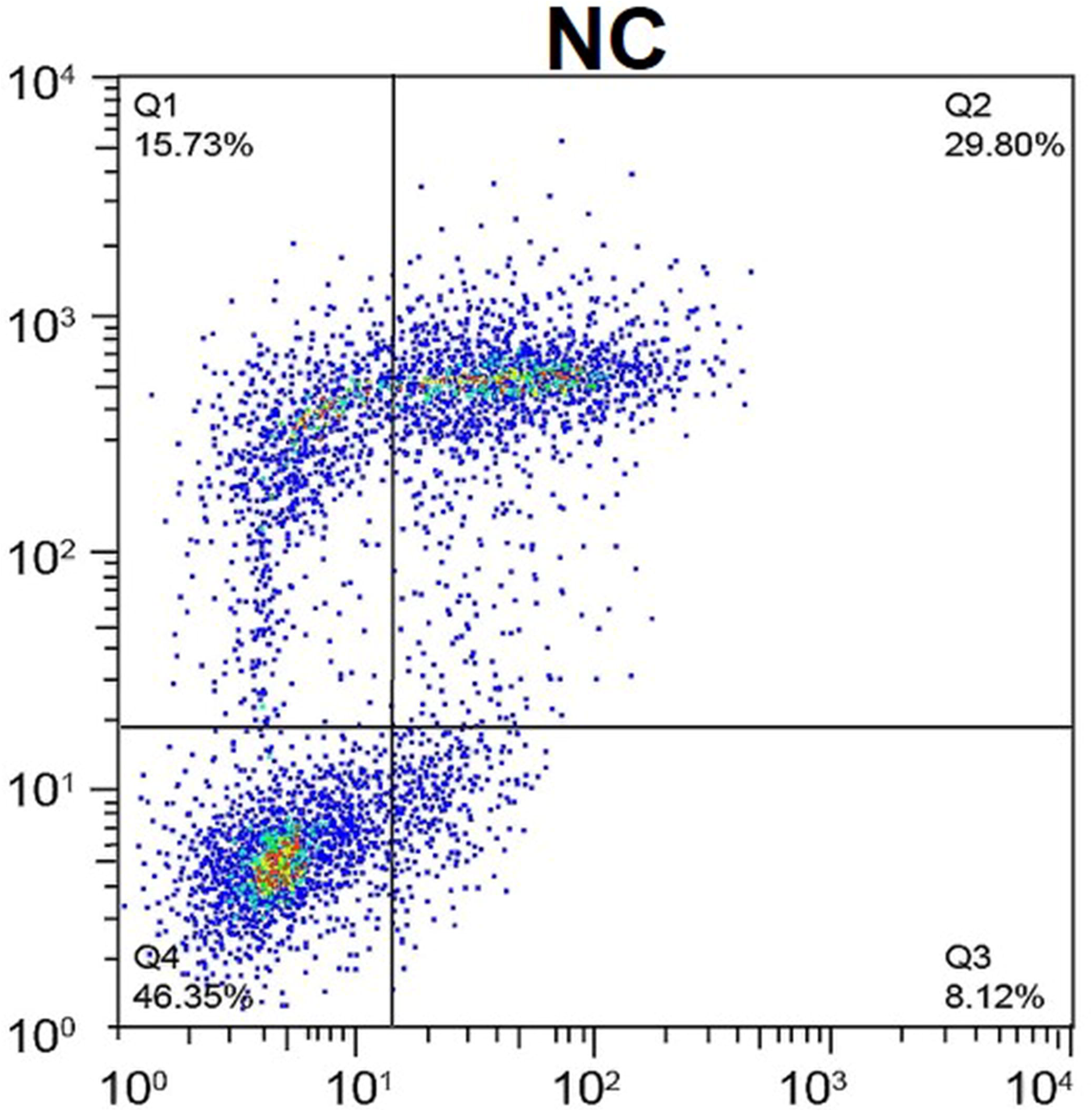
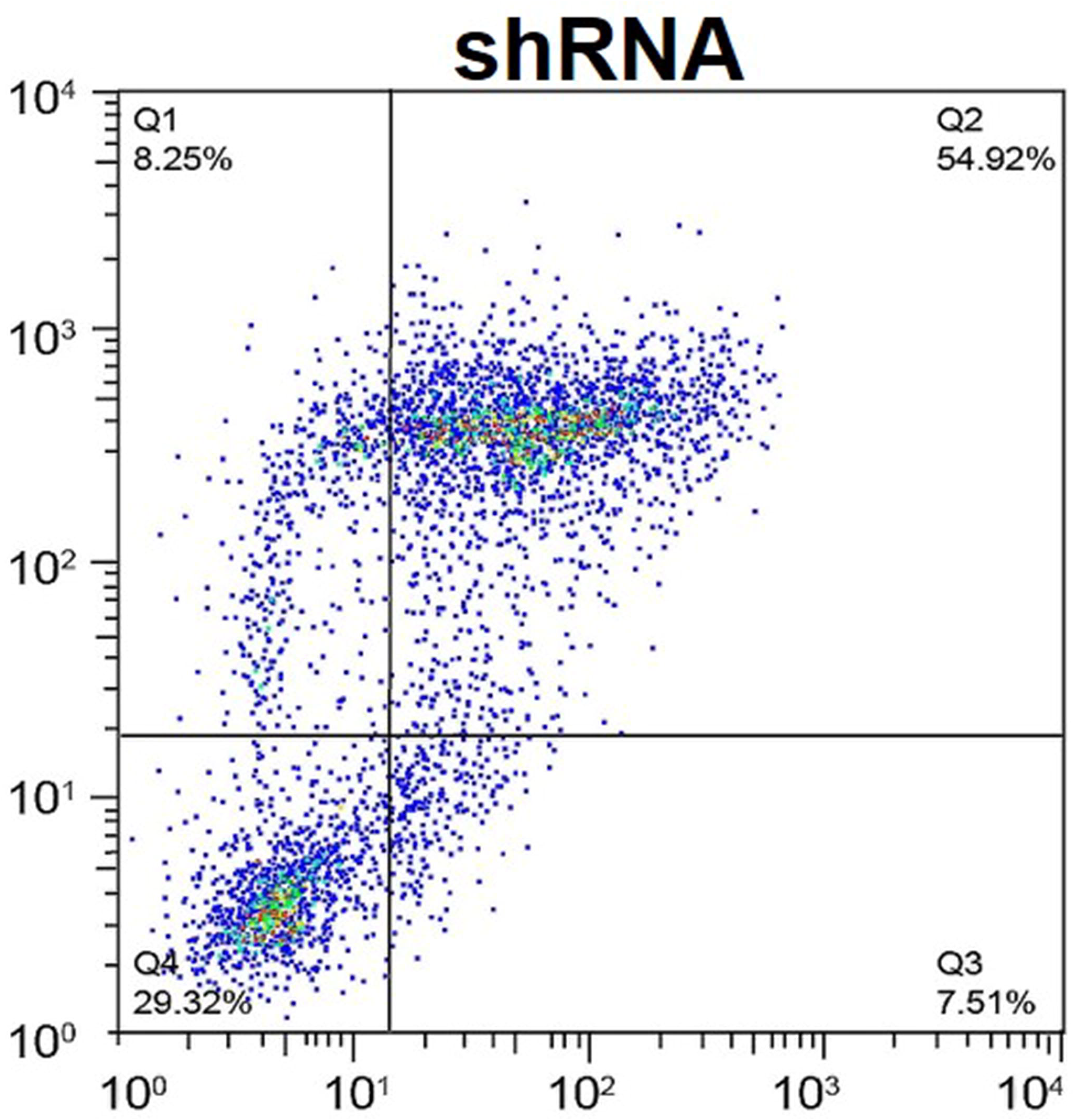
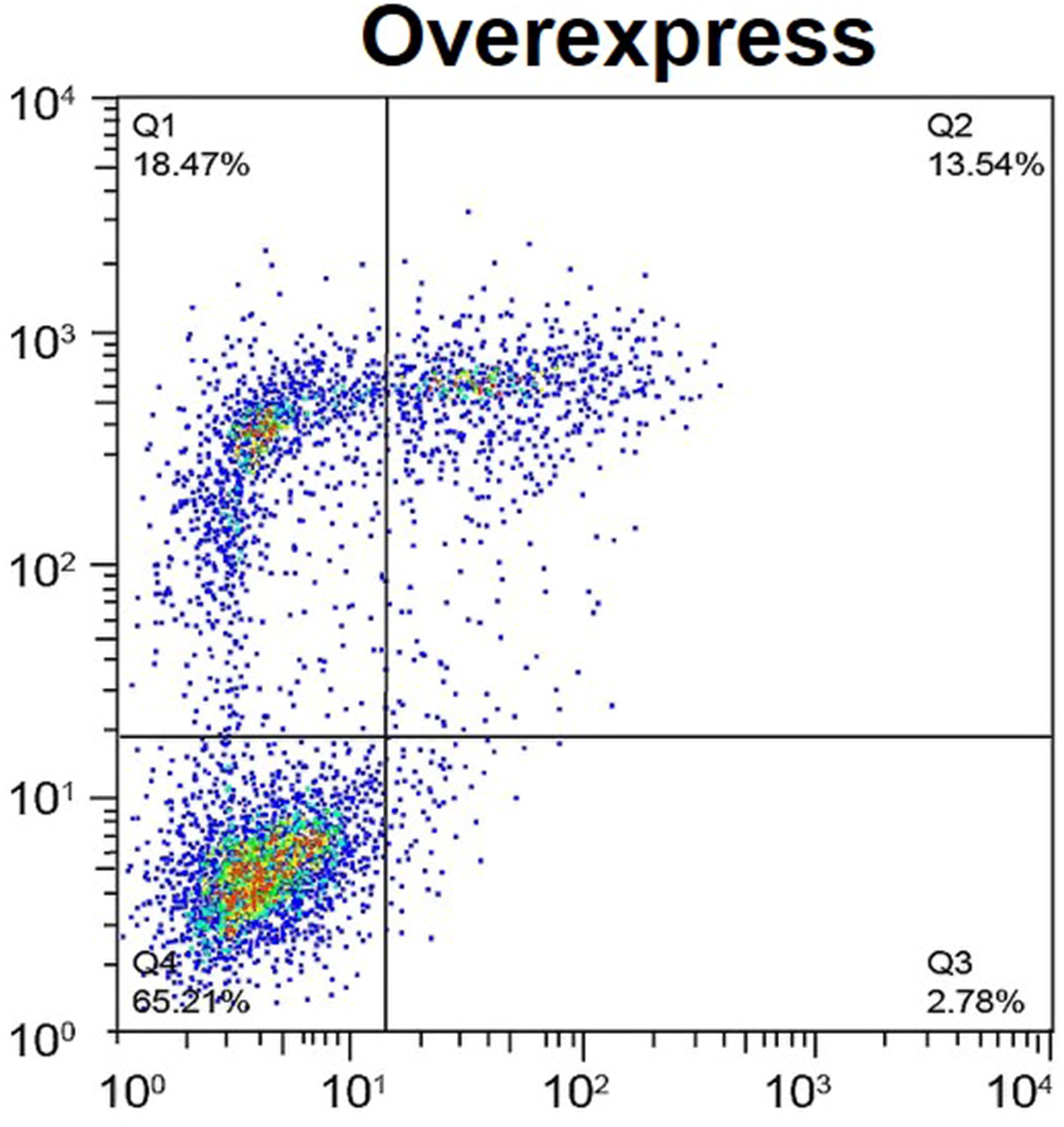
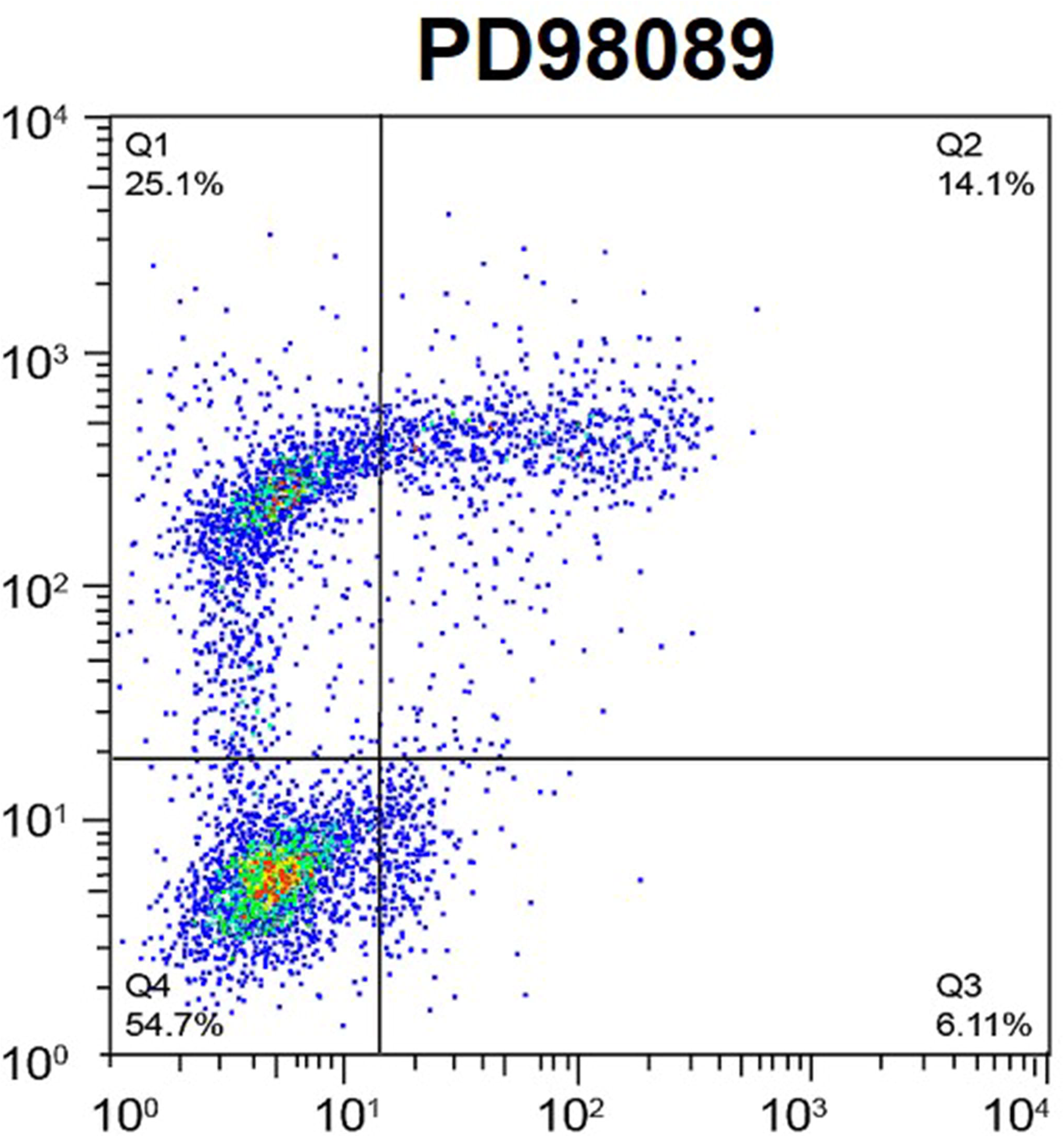
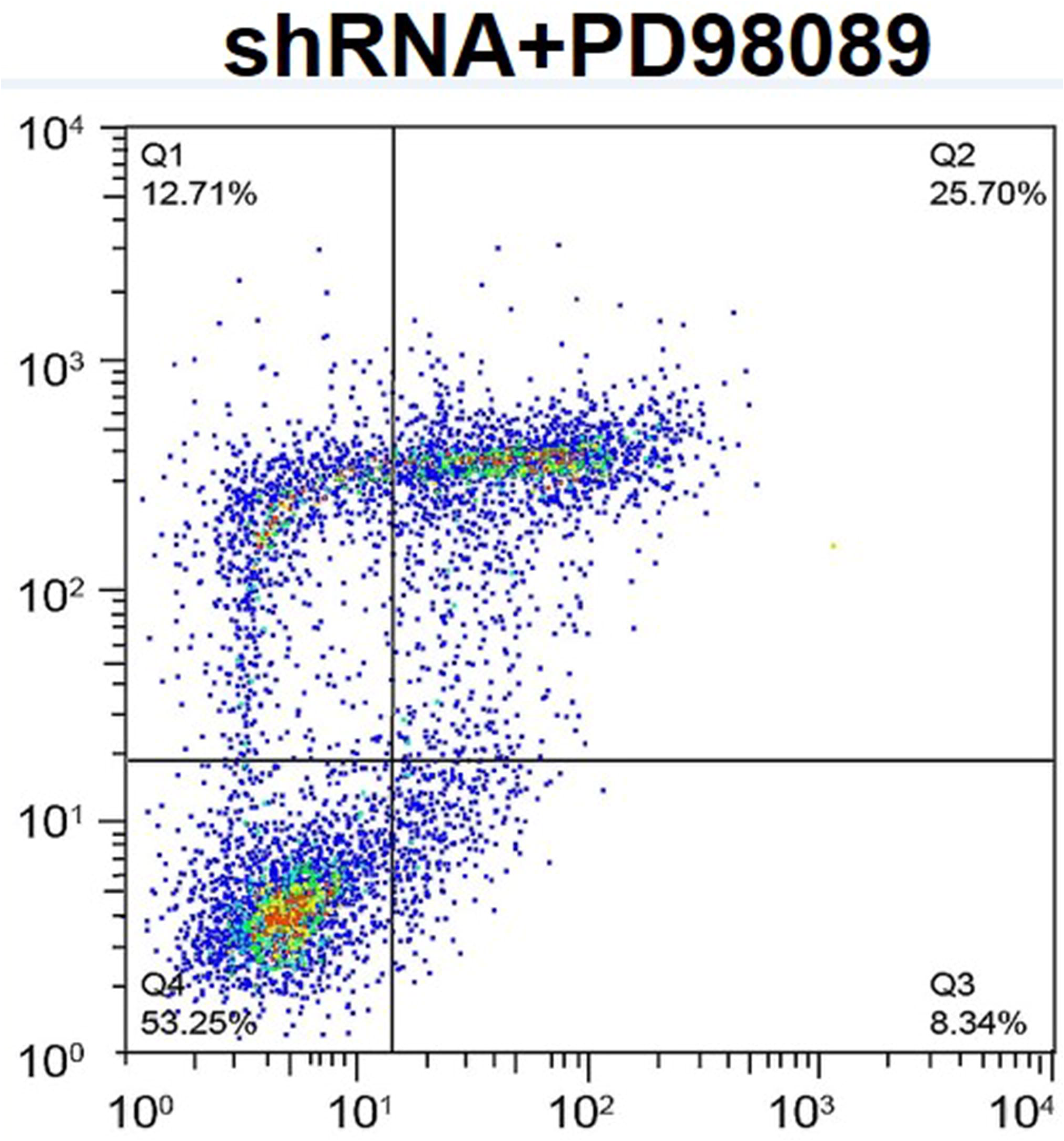
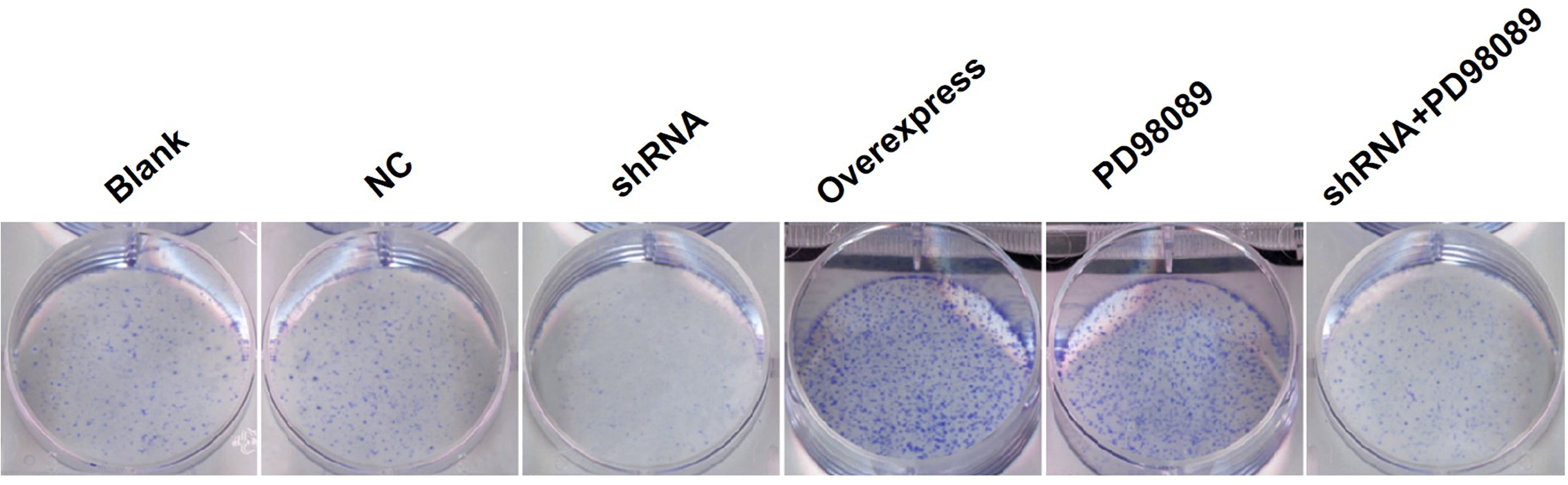
Analysis of cell proliferation and apoptosis. (A) An MTT assay showed that IGFBP5 overexpression increased cellular proliferation at day 10. (B-N) Flow cytometry and colony formation assays demonstrated that IGFBP5 overexpression could effectively promote NP cell proliferation and inhibit NP cell apoptosis when compared with IGFBP5-shRNA. NP=nucleus pulposus.

### IGFBP5 exerts its functions by inhibiting the ERK signaling pathway

The NP cells in each group were transfected and showed high transfection efficiency (Fig. 3A). To explore whether IGFBP5 exerts its function through the ERK signaling pathway, which contributes to NP cell proliferation and survival, we performed RT-PCR and Western blot analyses to examine the levels of genes and related proteins, including ERK, pERK, Bax, caspase-3 and Bcl2. The expression of ERK, pERK, Bax and caspase-3 were decreased, while the level of Bcl2 was increased in NP cells that stably overexpressed IGFBP5 mRNA (Fig. 3B and 3C). Additionally, the effect of IGFBP5 mRNA overexpression on NP cells was similar to those induced by the ERK inhibitor, PD98059. In contrast, the expression of ERK, pERK, Bax and caspase-3 was significantly upregulated, while the level of Bcl2 was downregulated following transfection with IGFBP5-shRNA (Fig. 3B and 3C). No differences in the expression levels of these genes and related proteins were observed between the IGFBP5-shRNA + PD98059, blank and NC groups (all Ps > 0.05). These data indicate that IGFBP5 mRNA inhibits the ERK signaling pathway in IDD.

**Fig. 3.**
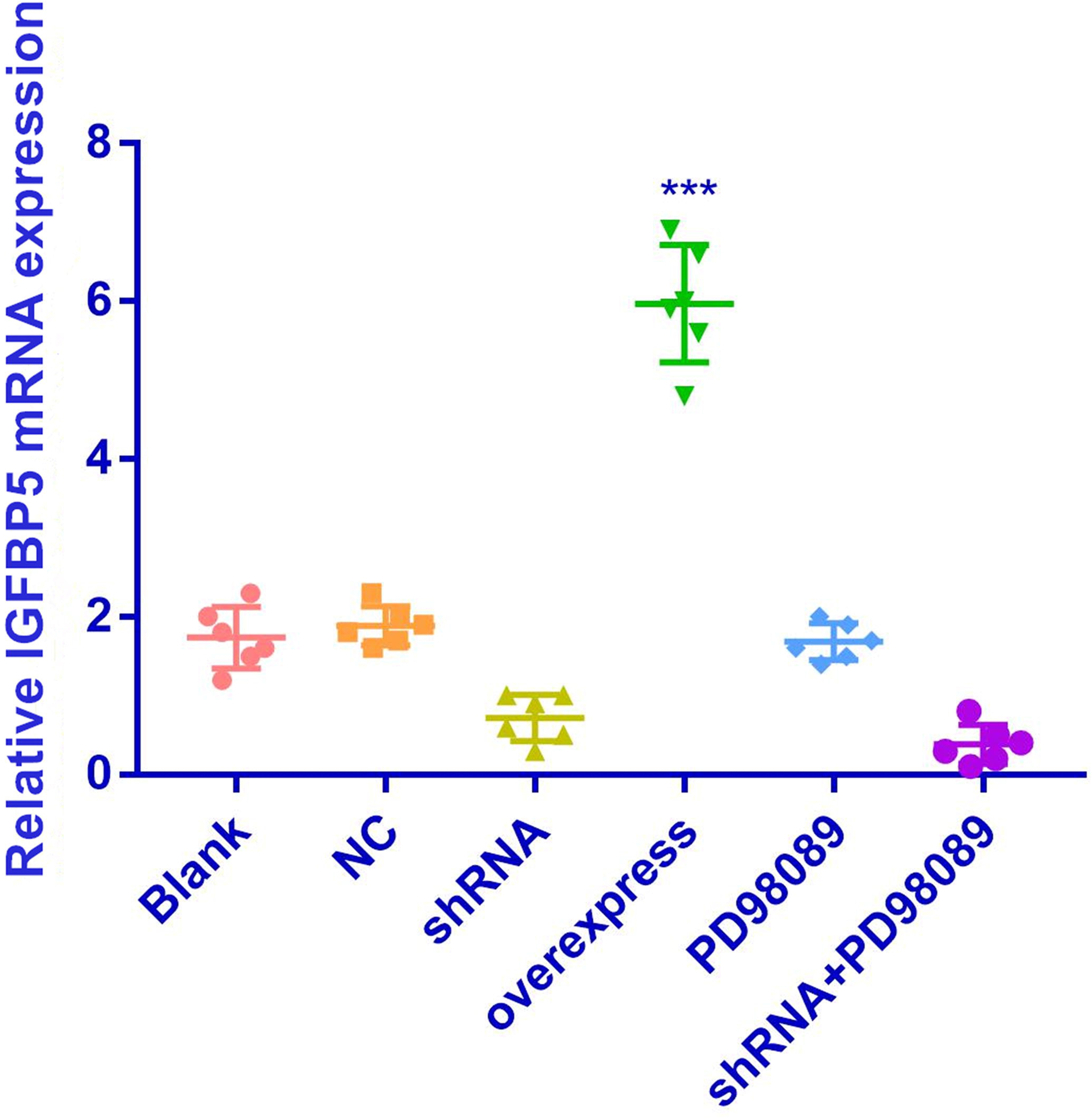
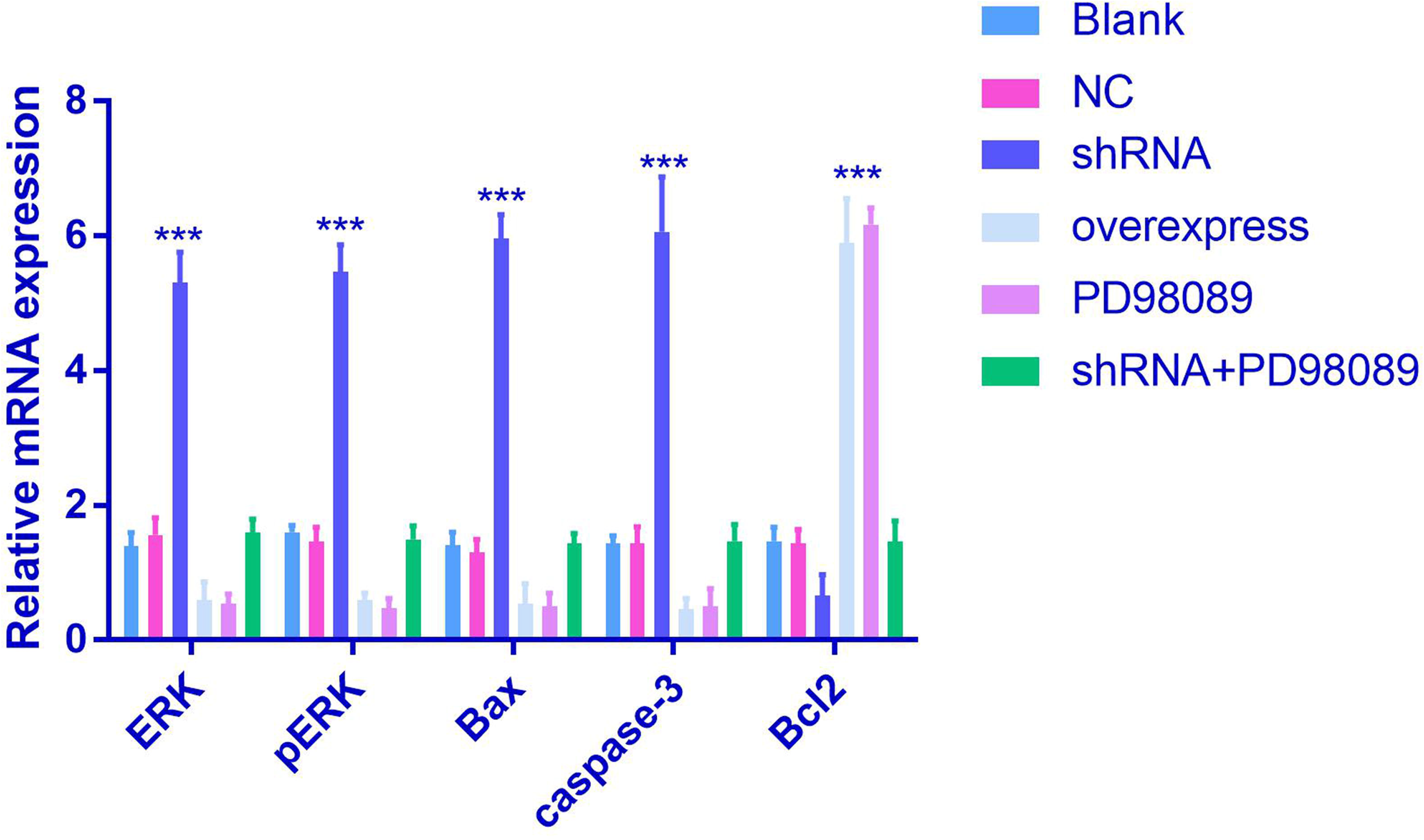
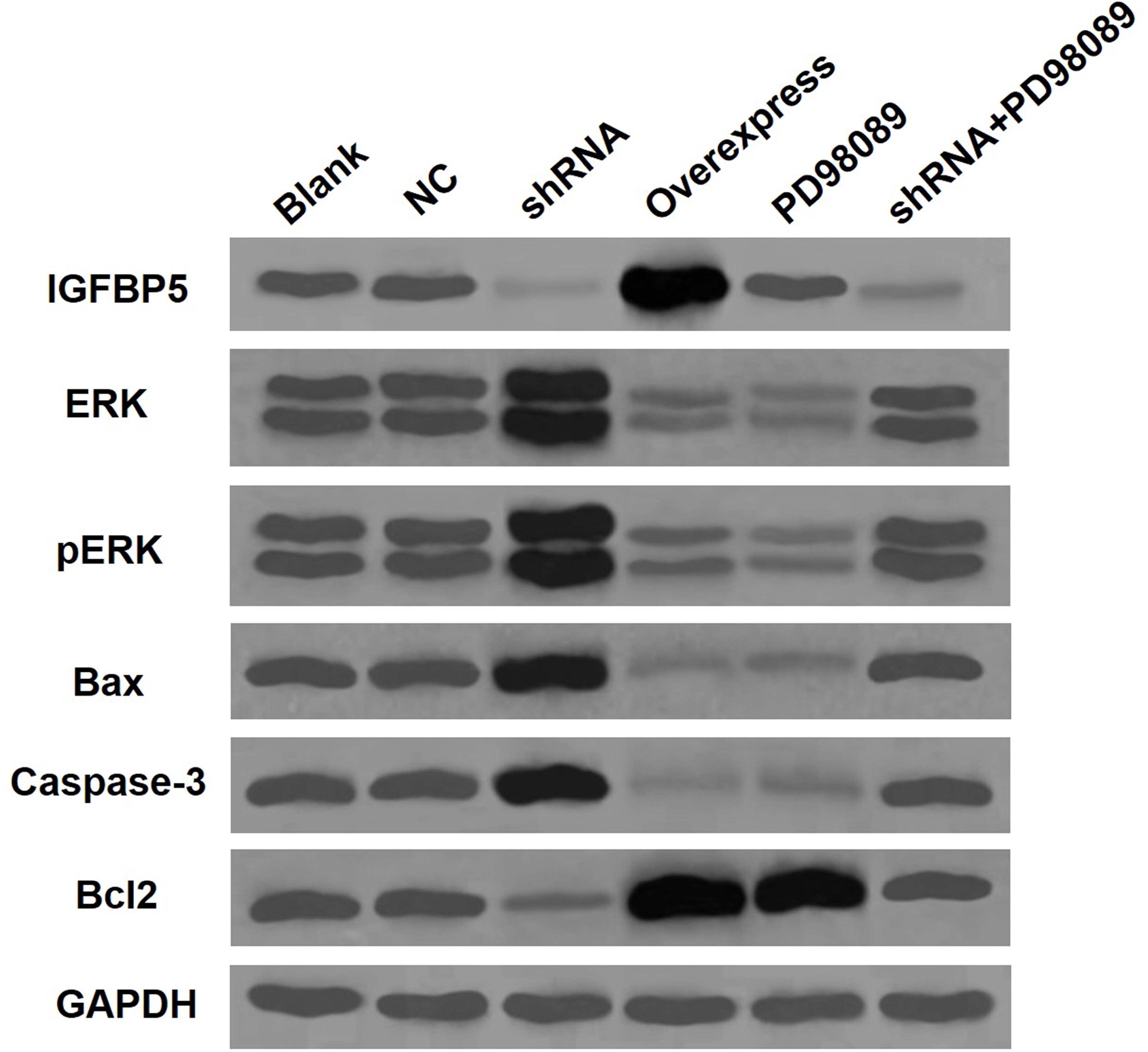
IGFBP5 exerts its function by inhibiting the ERK signaling pathway. (A, B) Overexpression of IGFBP5 induced effects on NP cells similar to those induced by PD98089. In contrast, knockdown of IGFBP5 abrogated the effects on NP cells, followed by ERK and pERK upregulation. (B, C) In NP cells treated with IGFBP5-shRNA, the expression levels of ERK, pERK, Bax and caspase-3 were significantly increased, while the level of Bcl2 was decreased compared with the control. Values presented as the mean±SD. ***p < 0.001. NP=nucleus pulposus.

### IGFBP5 inhibited disc degeneration in the IDD model

To investigate the role of IGFBP5 in vivo, a rat model of IDD was generated. As shown in Fig. 4A and 4b, the progression of IDD in the IDD + Igfbp5 group was inhibited compared with the IDD group. Moreover, the IGFBP5 level in NP cells was lower in IDD rats when compared with IDD + Igfbp5 rats (P<0.0001), which was confirmed by immunofluorescence staining (Fig. 4C and 4D). The levels of ERK, pERK, Bax, caspase-3 and Bcl2 were measured by RT-qPCR and Western blotting (Fig. 4D and 4E). Compared with IDD rats, the ERK, pERK, Bax and caspase-3 expression levels were lower in IDD + Igfbp5 rats (P< 0.0001). In contrast, the level of Bcl2 was higher in IDD +Igfbp5 rats than in IDD rats (P< 0.0001).

**Fig. 4.**
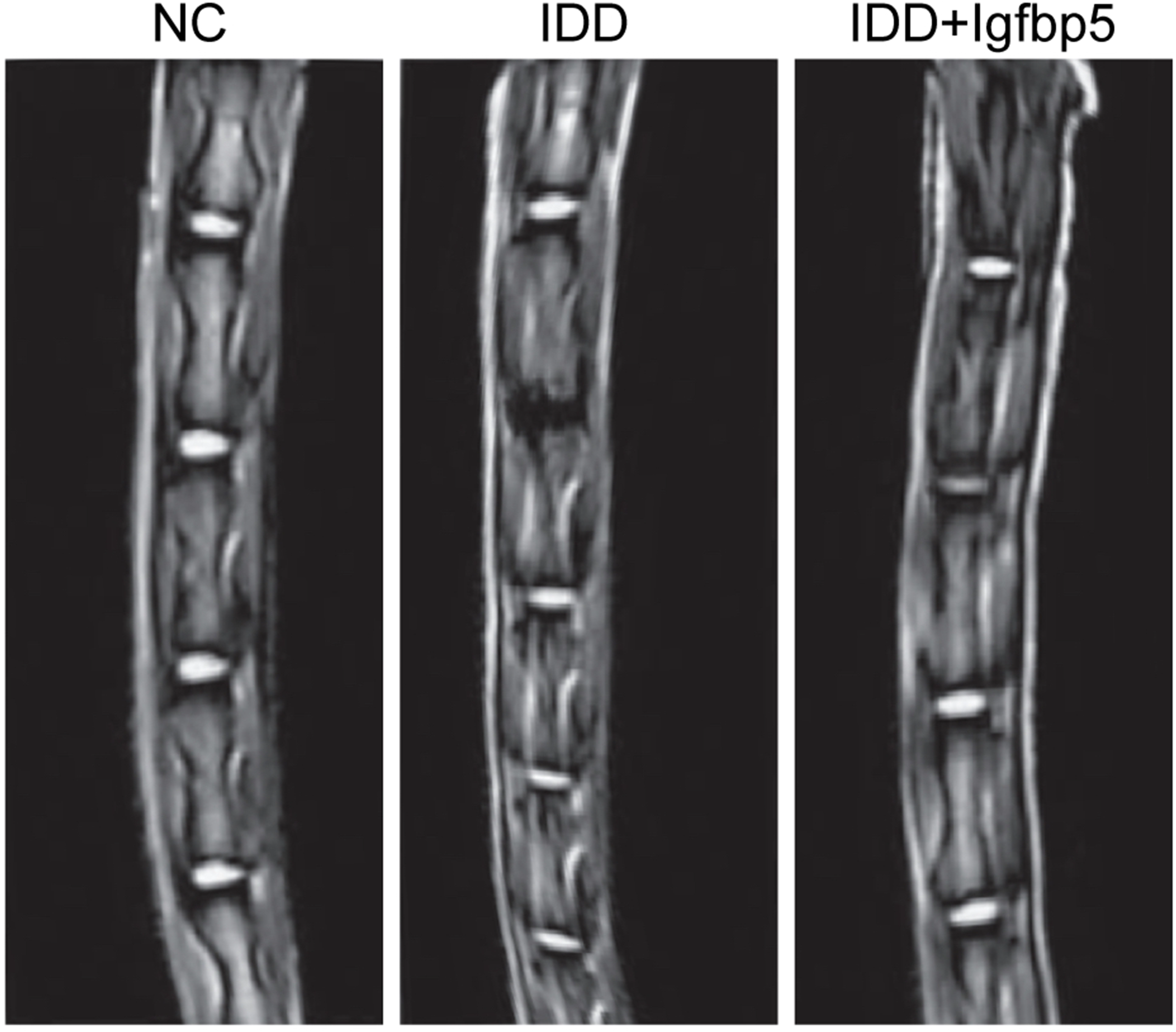
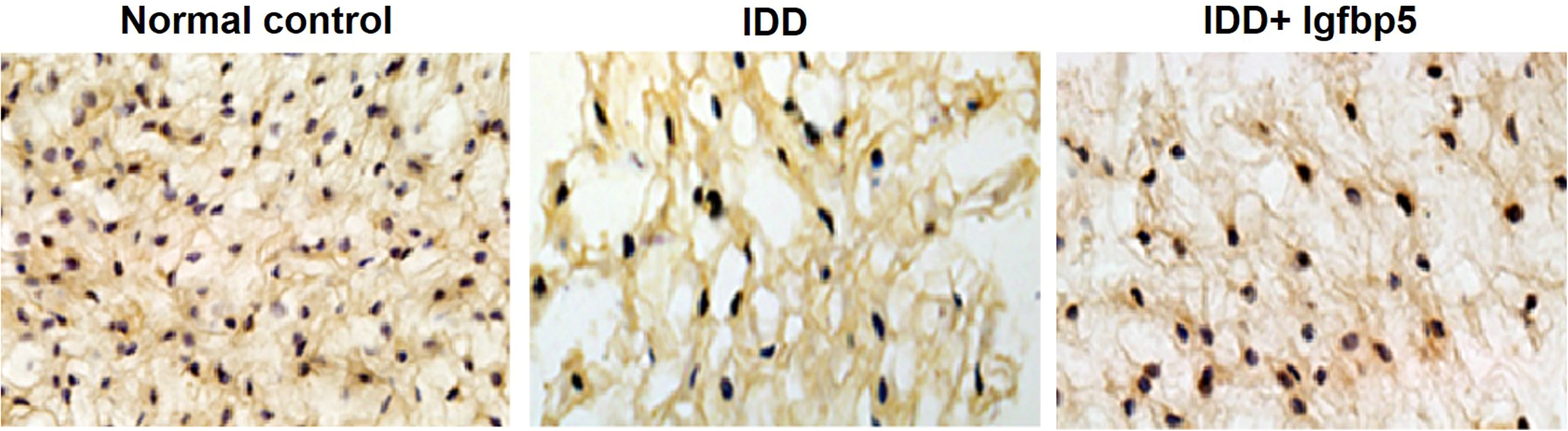
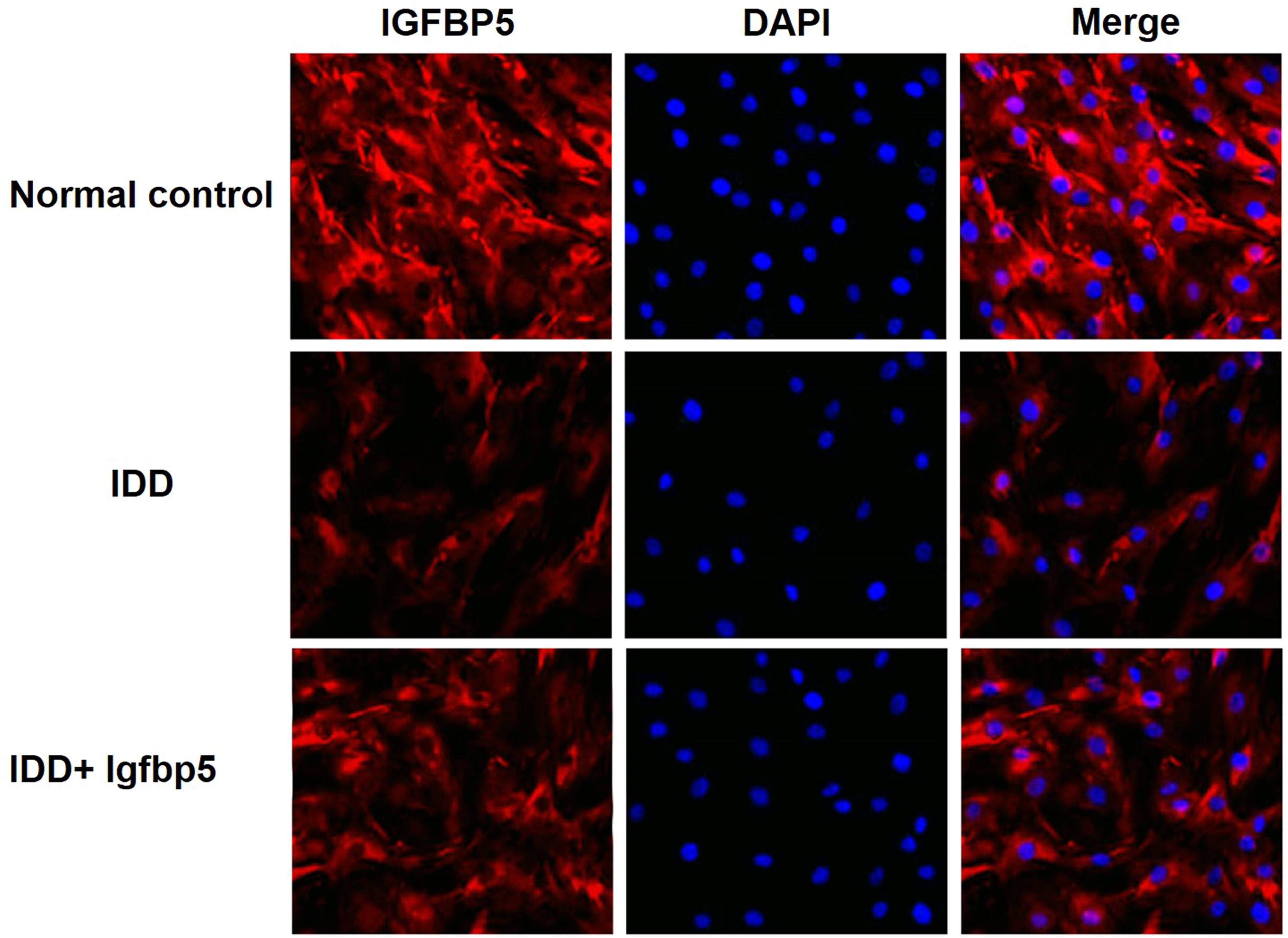
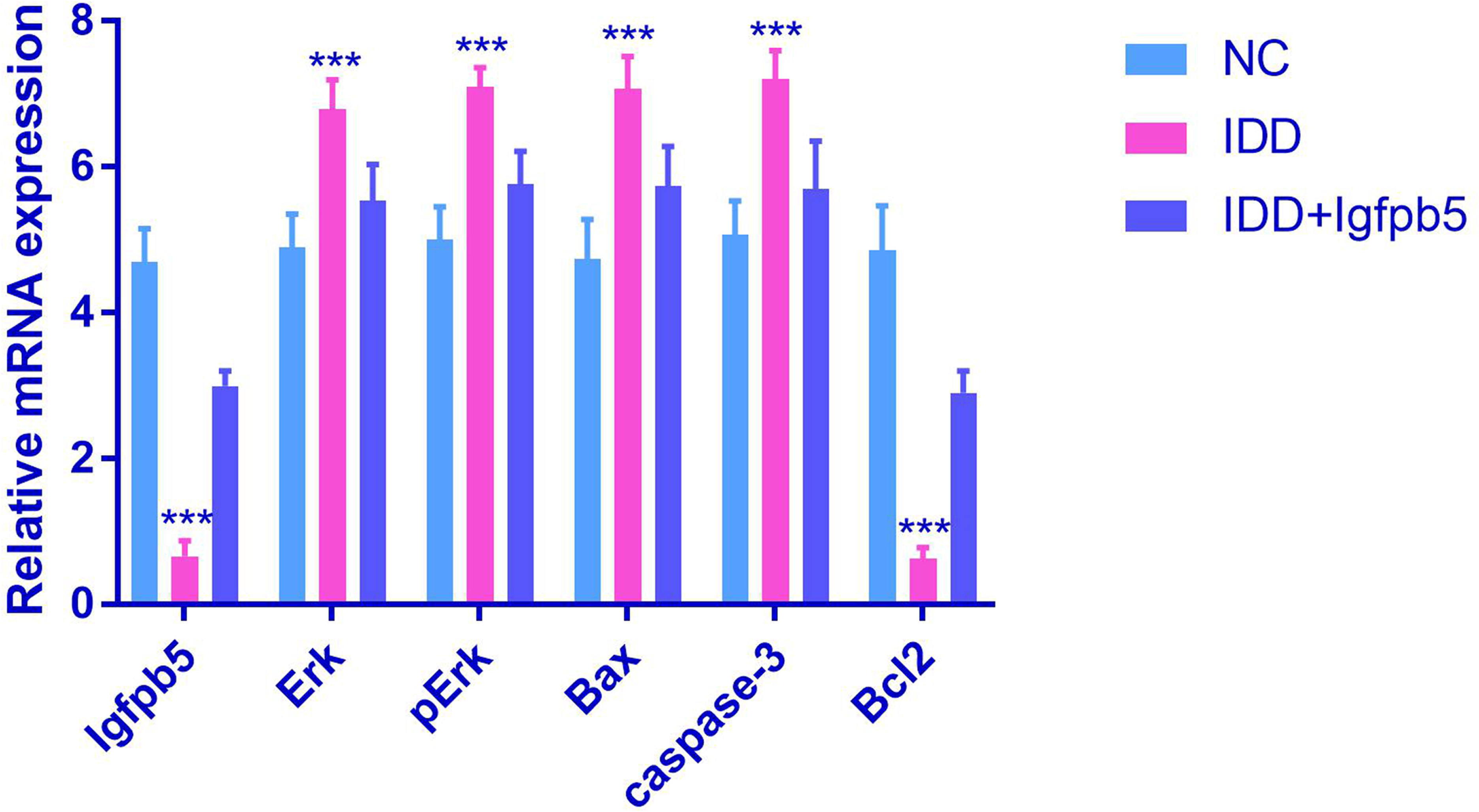
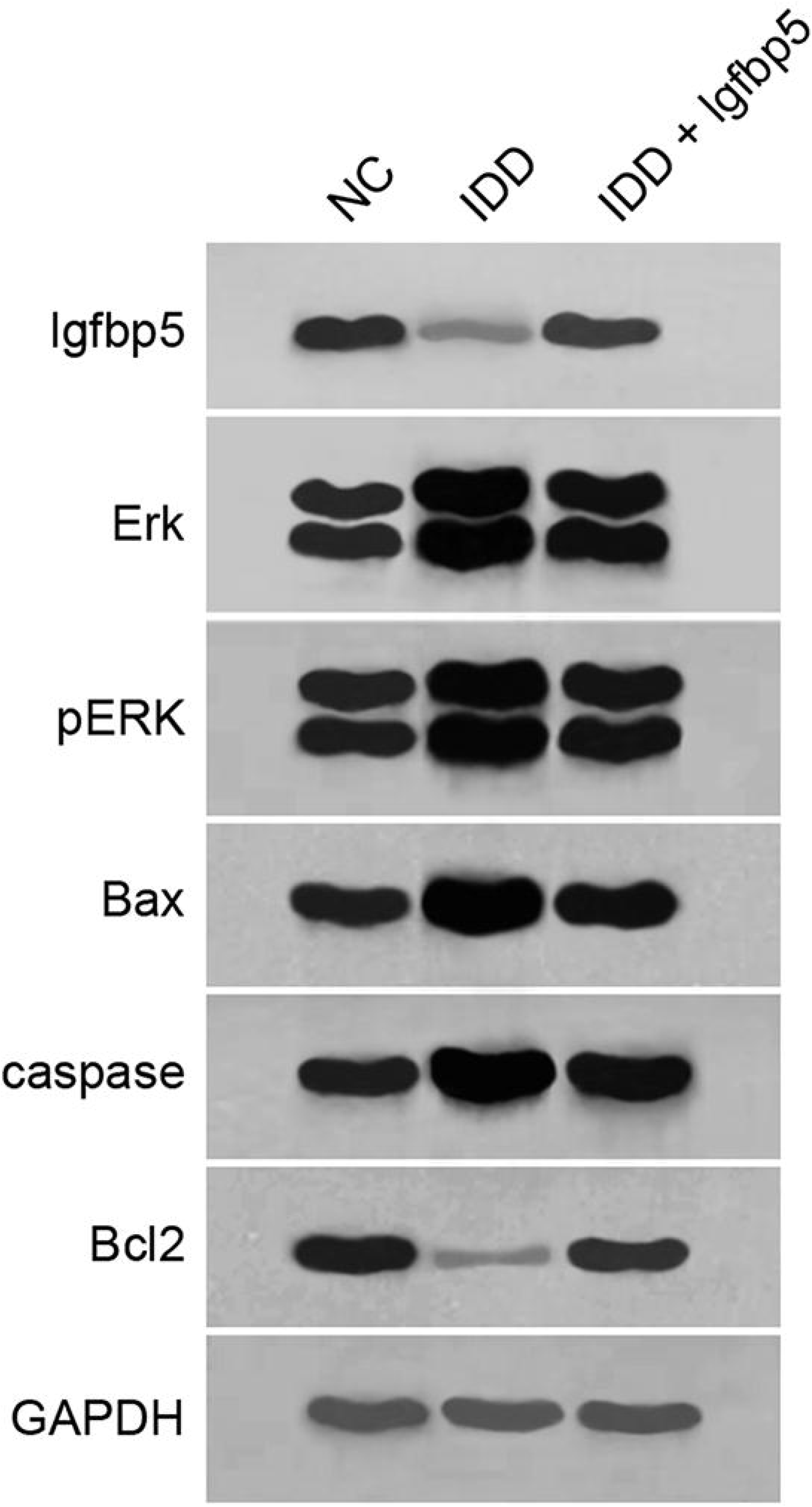
Effect of IGFBP5 in IDD rats. (A, B) Severe intervertebral disc degeneration was induced in the rats in the IDD group. Igfbp5 mRNA expressing treatment abrogated the effects on the intervertebral disc tissues. (C, D) The expression level of IGFBP5 in NP cells was detected by RT-qPCR and immunofluorescence staining. (C, E) The levels of ERK, pERK, Bax, caspase-3 and Bcl2 were measured by RT-qPCR and Western blotting. Data are shown as the mean±SD of five rats in each group. NC=normal control group, IDD=IDD model group, IDD + Igfbp5=IDD model rats treated with the Igfbp5 mRNA expressing lentivirus. ***P<0.001.

## Discussion

IGFBP5, a key member of the IGF axis, has been known to play important roles in diverse biological and pathological processes, including cell proliferation, differentiation, apoptosis and carcinogenesis (Beattie et al., 2006; Karabulut et al., 2016; Sanada et al., 2016). In this study, we demonstrated that dysregulated IGFBP5 expression induced NP cell apoptosis. Mechanistically, the downregulation of IGFBP5 led to activated ERK, which induced the expression of gene products with death-promoting activity. Although regulation of the ERK pathway by IGFBP5 has been reported in human intestinal smooth muscle cells and periodontal ligament stem cells (Kuemmerle and Zhou, 2002; Wang et al., 2016), this molecular pathway has not been reported in nuclear pulposus cells in relation to IDD. Here, we provide evidence for a mechanistic link between IGFBP5 and the ERK signaling pathway in NP cells during IDD. Our novel findings suggest that downregulation of IGFBP5 might participate in the development of IDD.

In the current study, we first performed state-of-the-art NGS in degenerative NP tissues vs. normal disc tissues to identify IDD-specific mRNAs. Using the Illumina HiSeq 2500 platform and subsequent qRT-PCR, IGFBP5 mRNA was found to be significantly downregulated in patients compared with corresponding controls, and thus was selected for further analysis. Moreover, the IGFBP5 expression level was significantly lower in degenerative NP tissues and was significantly correlated with the disc degeneration grade. To further investigate the role of IGFBP5 in the pathogenesis of IDD, we thereby performed functional analysis of IGFBP5 to investigate the relationship between IGFBP5 and NP cell proliferation and apoptosis. In our study, NP cell proliferation was significantly increased by the overexpression of IGFBP5, whereas it was significantly inhibited by the decreased expression of IGFBP5 through transfection with IGFBP5-shRNA. In addition, the overexpression of IGFBP5 led to an increase in cell growth and the percentage of cells in the S phase of the cell cycle, together with a decrease in the cell apoptosis rate and the percentage of cells in the G1 phase. Contrasting results were observed following the transfection with IGFBP5-shRNA. These findings suggest that increased NP cell proliferation induced by overexpression of IGFBP5 may be one of the possible mechanisms of IDD development.

The regulation of cell proliferation and apoptosis is a complex process and is associated with several signaling pathways (Masuda, 2008; Masuda and An, 2006; Pratsinis and Kletsas, 2007; Wang et al., 1999; Zhang and Liu, 2002). Extracellular signal-regulated kinase (ERK), which belongs to the family of mitogen-activated protein kinases (MAPKs), exerts an important role on the regulation of cell differentiation (Li et al., 2014; Sparta et al., 2015). It is activated through a sequential phosphorylation cascade that amplifies and transduces signals from the cell membrane to the nucleus (Caunt and McArdle, 2012; Iwamoto et al., 2016; Maryu et al., 2016; Shindo et al., 2016). Both in vitro and in vivo data have indicated that the proliferation and apoptosis of human IVD cells is regulated by exogenous and autocrine growth factors mainly via the ERK pathway (Pratsinis et al., 2012; Pratsinis and Kletsas, 2007; Uchiyama et al., 2007). To determine the potential link between IGFBP5 and the ERK signaling pathway, IGFBP5 mRNA was overexpressed in NP cells by transfection and the results showed that the expression of both ERK and phosphorylated ERK were attenuated in IGFBP5-overexpressing cells, indicating that IGFBP5 inhibits cell apoptosis by regulating the ERK pathway. Conversely, the phosphorylation of these kinases was enhanced in IGFBP5 knockdown cells, which was confirmed by Western blot and RT-PCR analyses. In addition, the activity of ERK pathway associated proteins, such as Bax (Iwayama and Ueda, 2013; Ma et al., 2014; Yating et al., 2015) and caspase-3 (Sudo and Minami, 2011; Wang et al., 2011), and the downregulation of antiapoptotic proteins, such as Bcl2 (Mebratu and Tesfaigzi, 2009; Roskoski, 2012; Sudo and Minami, 2010), is associated with the upregulation of apoptosis execution. Our results obtained from the qRT-PCR and Western blot analyses indicated that the Bax and caspase-3 levels were significantly increased, while the level of Bcl2 was significantly decreased in the IGFBP5-shRNA group. All of the above findings demonstrate that the silencing of IGFBP5 induced the apoptosis of NP cells by activating the ERK pathway, which further upregulated the Bax and caspase-3 levels and downregulated the Bcl2 level.

Another interesting aspect of our study is that we, for the first time, demonstrated the role of IGFBP5 in IDD using a rat model. Our data revealed that the IGFBP5 mRNA level in NP cells was lower in IDD rats when compared with IDD + Igfbp5 rats (P<0.0001). Additionally, the Bcl2 expression level was higher in IDD + Igfbp5 rats, while the levels of ERK, pERK, Bax and caspase-3 were higher in IDD rats. Collectively, our findings provide novel mechanistic insight that was previously unrecognized and highlight that the suppression of IGFBP5 is an early phenomenon in IDD that may trigger the initiation of IDD.

Taken together, this study provides the discovery and validation of IDD-specific mRNA transcriptome profiles generated by NGS. We identified that IGFBP5 was downregulated in human degenerative NP tissues and that its level was associated with the disc degeneration grade. In addition, the inhibitory effects of downregulated IGFBP5 are, in part, mediated through the ERK signaling pathway. Using in vivo and in vitro studies, functional characterization of the role of IGFBP5 revealed that the loss of its expression is an event in IDD pathogenesis. Therefore, strategies to maintain the expression or to prevent the repression of IGFBP5 have the potential to become a possible therapeutic and/or preventive approach for degenerative disc disease.

## Material and methods

### Patient samples

Between February 2015 and March 2017, a total of 129 lumbar NP specimens were obtained from 129 patients with degenerative disc disease undergoing discectomy (mean age: 54.7±9.6, range: 44–69 years). Of them, 10 NP specimens were randomly selected for mRNA Solexa sequencing. The surgical indications were as follows: (1) failed conservative treatment and (2) progressive neurologic deficits such as progressive motor weakness or cauda equine syndrome. Patients with isthmic or degenerative spondylolisthesis, lumbar stenosis, ankylosing spondylitis, or diffuse idiopathic skeletal hyperostosis were excluded. The control samples were taken from 112 patients with fresh traumatic lumbar fracture (mean age: 21.1±3.4, range: 18–23 years) who underwent anterior decompressive surgery due to neurological deficits. Of them, 10 NP specimens were randomly selected for mRNA Solexa sequencing. In an attempt to reduce the differences between the individuals and to obtain more data, a 10 (pathological) vs. 10 (control) experimental design was thus employed for the mRNA Solexa sequencing in our study.

Routine MRI scans of the lumbar spine were taken of these patients before surgery and the degree of disc degeneration was graded from the T2-weighted images using the Pfirrmann classification (Pfirrmann et al., 2001). According to the modified classification system of the International Society for the Study of the Lumbar Spine (Williams et al., 1982), 101 samples were protrusions, 21 were sequestration, and 7 were subligamentous extrusions. Tissue specimens were first washed with phosphate-buffered saline (PBS). Thirty-eight of the 129 samples were obtained from the L4/L5 level, whereas 91 were from L5/S1. The study protocol was approved by the Ethics Committee of our institution, and written informed consent was obtained from each participant.

### Isolation and culture of human NP cells

Tissues specimens were first washed thrice with PBS. The NP was then separated from the annulus fibrosus using a stereotaxic microscope and cut into pieces (2 to 3 mm^3^). The samples were digested by 0.25 mg/mL type II collagenase (Invitrogen, Carlsbad, CA, USA) for 12 h at 37°C in Dulbecco’s modified Eagle medium (DMEM; GIBCO, Grand Island, NY, USA) until the tissue blocks disappeared. After isolation, NP cells were resuspended in DMEM containing 10 % FBS (GIBCO, NY, USA), 100mg/ml streptomycin, 100 U/ml penicillin, and 1 % L-glutamine, and then incubated at 37 °C in a humidified atmosphere with 95 % air and 5 % CO_2_. The confluent cells were detached by trypsinization and seeded into 35-mm tissue culture dishes in complete culture medium (DMEM supplemented with 10 % FBS, 100 mg/ml streptomycin and 100 U/ml penicillin) in a 37 °C, 5 % CO_2_ environment. The medium was changed every 2 days. The second passage was used for the subsequent experiments.

### Cell transfections

The isolated cells were divided into the following groups: the blank group, the negative control (NC) group, the IGFBP5-shRNA group, the IGFBP5 overexpression group, the ERK pathway inhibitor group (PD98059), and the IGFBP5-shRNA + PD98059 group. Cells were seeded into 25-cm^2^ culture flasks and allowed to reach 30%-50% confluency. Serum-free medium (SFM, 100 µL) was used to dilute 5 µL of Lipofectamine 2000 (Invitrogen Grand Island, NY, USA) in sterile epoxy resin (EP) tubes, and then they were incubated at room temperature for 5 min. For the IGFBP5-shRNA group, short hairpin RNAs (shRNA) with the complementary sequences of the target genes were subcloned into the pLKO.1 lentiviral vector (Addgene, Cambridge, MA, USA). Viral packaging was prepared according to the manufacturer’s protocol (Clontech Laboratories, Addgene). For the viral infections, NP cells were plated overnight and then infected with retroviruses or lentiviruses in the presence of polybrene (6 μg/mL; Sigma-Aldrich, St. Louis, MO, USA) for 6 h. After 48 h, infected cells were selected with different antibiotics. Scrambled shRNA (Scramsh) was used as a control and was purchased from Addgene. The target sequence for the IGFBP5 shRNA (IGFBP5sh) was as follows: 5’-GCAGATCTGTGAATATGAA-3’.

For the IGFBP5 overexpression group, IGFBP5 was expressed in vitro using the gateway vector pLenti6.2/V5-DEST (Invitrogen, Grand Island, NY, USA), hereby referred to as pLenti6.2-IGFBP5. First, IGFBP5 was amplified from pCR4-TOPO-IGFBP5 (Thermo Scientific, Pittsburgh, PA, USA). Following the manufacturer’s protocols, IGFBP5 was then shuttled into the pENTR/D-TOPO vector (Invitrogen) and recombined into pLenti6.2/V5-DEST (Invitrogen). In addition, to suppress ERK expression, cells were treated with PD98059 according to the manufacturer’s instructions.

### RNA isolation

TRIzol (Invitrogen) and the RNeasy mini kit (Qiagen, Valencia, CA, USA) were applied for the extraction of the total RNA from NP cells according to the manufacturers’ protocols. Subsequently, RNA was eluted in 50 mL of nuclease-free water and stored at -80°C for further analysis. The RNA concentration was measured using a NanoDrop ND-1000 spectrophotometer (NanoDrop Technologies, Wilmington, DE, USA).

### Transcriptome sequencing

The total RNA of 10 patients and 10 controls was purified directly for transcriptome sequencing analysis using the Illumina HiSeq 2500 platform (San Diego, CA, USA) according to the manufacturer’s instructions. The library for sequencing was generated using an Illumina TruSeq RNA Sample Preparation Kit (Illumina, San Diego, CA, USA), which included procedures for RNA fragmentation, random hexamer-primed first strand cDNA synthesis, dUTP-based second strand cDNA synthesis, end-repair, A-tailing, adaptor ligation, and library PCR amplification. Image files were generated by the sequencer and processed to produce digital-quality data. After masking the adaptor sequences and removing the contaminated reads, clean reads were processed for in silico analysis. Then, a significance analysis of microarray (SAM) (version 4.0) was performed to select the mRNAs (Stanford University). Hierarchical cluster analysis was conducted using Gene Cluster 3.0 software (Stanford University).

### Establishment of the rat IDD model

Three-month-old Sprague–Dawley rats were used. All of the rats were of similar weight (approximately 380 g) to ensure that the tail discs at the position chosen for the experiment were of similar size such that when punctured with a 20-gauge needle, injuries of similar severity would be produced. Fifteen rats were randomly divided into 3 groups: normal control group (NC; n=5), IDD model group (IDD; n=5), and IDD model with the treatment of the Igfbp5 mRNA expressing lentivirus group (IDD + Igfbp5; n=5). Lenti-Igfbp5-Control and lenti-Igfbp5 lentiviral particles were produced by triple transfection of 293T cells (Invitrogen, Carlsbad, USA) with the vectors pLVX-Igfbp5-Control and pLVX-Igfbp5, respectively, along with psPAX2 and pMD2.G. All rats were anesthetized by intraperitoneal injection of chloral hydrate (250 mg/kg). Subsequently, the IDD + Igfbp5 and IDD groups underwent annulus fibrosus needle puncture to induce surgical IDD. Tail disc injection of 10 µL lentivirus was performed at 7 days post IDD surgery. The IDD groups accepted 10 µL lenti-NC, while the experimental group was injected with lenti-Igfbp5. Rats from each group were sacrificed at 8 weeks post-IDD surgery, and then the tails were dissected and processed for further histological evaluation. All animal experiments were approved by the Ethics Committees of our institution.

### Reverse transcription and quantitative real-time PCR

To determine the mRNA levels in the NP samples, reverse transcription (RT) and quantitative real-time PCR (qPCR) kits (Applied Biosystems) were used. RT reactions were performed using the RevertAid RT Reverse Transcription Kit (Thermo Scientific™) in a final volume of 12 μL (65°C for 5 min, ice bath for 2 min, 42°C for 60 min, 70°C for 5 min, and hold at -20°C).

Real-time PCR was performed using a standard TaqMan PCR protocol. The 20-μL PCR reactions included 4 μL diluted RT product, 20 μL TaqMan Universal PCR Master Mix with no AmpErase UNG, 1 μL TaqMan primers, and 5 μL nuclease-free water. The PCR primers were as follows: IGFBP5 (forward, ACCTGAGATGAGACAGGAGTC; reverse, GTAGAATCCTTTGCGGTCACAA), ERK (forward, ACTCACCTCTTCAGAACGAATTG; reverse, CCATCTTTGGAAGGTTCAGGTTG), pERK (forward, CCTGAACCTTCCAAAGATGGC; reverse, TTCACCAGGCAAGTCTCCTCA), Bax (forward, CCCGAGAGGTCTTTTTCCGAG; reverse, CCAGCCCATGATGGTTCTGAT), caspase-3 (forward, CTGAACAGCTCCGAGGAAAC; reverse, TGGATATTCAGCCCTTTTGG), Bcl2 (forward, GGTGGGGTCATGTGTGTGG; reverse, CGGTTCAGGTACTCAGTCATCC), and GAPDH (forward, GGAGCGAGATCCCTCCAAAAT; reverse, GGCTGTTGTCATACTTCTCATGG). All reactions were performed in triplicate on a 7500 Real-time system (Applied Biosystems, Foster City, CA, USA) with the following conditions: 95 °C for 10 min, followed by 40 cycles at 95 °C for 15 s, and 60 °C for 1 min. Glyceraldehyde-3-phosphate dehydrogenase (GAPDH) was used as the internal control. The relative quantitative method was used to analyze the data, applying the 2-ΔΔCt method to determine the relative expression of each target gene. The experiments were repeated three times independently.

### Western blotting

According to standard methods, Western blot analysis was performed. Briefly, proteins were separated on 10 % SDS-PAGE gels and then transferred to PVDF membranes (Amersham, Buckinghamshire, UK). Membranes were blocked using 5 % nonfat dried milk for 2 h and then incubated for 12 h with an anti-IGFBP5 antibody (1:1000; Abcam, Cambridge, UK), anti-ERK antibody (1:1000; Abcam, Cambridge, UK), anti-pERK antibody (1:500; Abcam, Cambridge, UK), anti-Bax antibody (1:1000; Abcam, Cambridge, UK), anti-caspase-3 antibody (1:1000; Abcam, Cambridge, UK), anti-Bcl2 antibody (1:1000; Abcam, Cambridge, UK), or anti-GAPDH antibody (1:10000; Abcam, Cambridge, UK). After washing in TBST (10 mM Tris, pH 8.0, 150 mM NaCl and 0.1% Tween 20), membranes were incubated for 2 h in goat anti-rabbit antibody (1:5000; Abcam, Cambridge, UK). Normalization was performed by blotting the same membranes with an antibody against GAPDH. The relative expression was quantified using Quantity One software, version 4.52 (Bio-Rad).

### 3-(4,5-Dimethylthiazol-2-yl)-2,5-diphenyltetrazolium bromide and colony formation assays

After the density of the transfected cells reached approximately 80%, cells were washed twice with PBS, digested in 0.25% pancreatin, and single-cell suspensions were obtained. After 7 and 14 days of culture in 24-well plates, cell proliferation was assessed by 3-(4,5-dimethylthiazol-2-yl)-2,5-diphenyltetrazolium bromide (MTT) assay. Briefly, the cells were cultured in 500 mL of MTT solution (250 mg in DMEM-HG) at 37°C for 4 h. During this incubation period, water-insoluble formazan crystals were formed. Then, the formazan crystals were dissolved by adding 300 mL of dimethyl sulfoxide. Absorbance was measured at a wavelength of 570 nm using a Spectra MAX microplate reader (Molecular Devices, Sunnyvale, CA, USA).

For the colony formation assay, NP cells at the logarithmic growth phase in each transfected group were selected and washed once with PBS. Then, NP cells in 2 ml of medium were plated per well in six-well plates. Following 10 days of incubation, the colonies were quantified after fixation with methyl alcohol and staining with hematoxylin. These experiments were performed three times.

### Flow cytometry

Apoptosis was evaluated by staining cells with both Annexin V-FITC and propidium iodide (PI), according to the manufacturer’s instructions. Annexin V-FITC was employed to quantitatively determine the percentage of cells undergoing apoptosis. It relies on the property of cells to lose membrane asymmetry in the early phase of apoptosis. In apoptotic cells, the membrane phospholipid phosphatidylserine is translocated from the inner leaflet of the plasma membrane to the outer leaflet, thereby exposing phosphatidylserine to the external environment. Cells that were positively stained with Annexin V-FITC and negatively stained for PI were considered apoptotic. Cells that were positively stained for both Annexin V-FITC and PI were considered necrotic. To quantify apoptosis, the cells were washed with cold PBS solution and then resuspended in binding buffer (10 mmol/L HEPES (N-2-hydroxyethylpiperazine-N‚-2-ethanesulphonic acid)/NaOH [pH 7.4], 140 mmol/L NaCl, and 2.5 mmol/L CaCl_2_). The cells were stained with 5 mL Annexin V-FITC and 10 mL PI and then analyzed with EpicsAltra (Beckman Coulter, CA, USA) flow cytometry (FCM).

### Immunohistochemical staining

Immunohistochemical staining was performed on intervertebral discs. Endogenous peroxidase was inactivated by incubating with 0.3 % H_2_O_2_ in PBS for 15 min, followed by a 30-min block step with PBST/1 % BSA. Sections were incubated with rabbit anti-collagen II (Abcam, Cambridge, UK) at 10 μg/ml in PBST and 1% BSA at 4°C overnight. Goat anti-rabbit IgG (H+L) (Abcam, Cambridge, UK) was used as the secondary antibody for collagen II staining at 2 µg/ml for 1 h at room temperature.

### Immunofluorescent staining

Coverslips were placed into 24-well plates where NP cells were plated for 48 h. The medium was removed and the cells were washed twice with PBS and then fixed with 3.5 % formaldehyde for 30 min at 37 °C. The cells were rinsed with PBS three times, permeabilized with 0.1 % Triton X-100 in PBS for 20 min, and blocked with 3 % BSA and 0.05 % Tween 20 in PBS for 30 min at room temperature. After blocking, cells were incubated with rabbit polyclonal anti-IGFBP5 (Abcam, Cambridge, UK) at 4 µg/mL overnight at 4 °C. The cells were then treated with fluorescent goat anti-rabbit secondary antibody (Abcam, Cambridge, UK) at 2 µg/ml for 1 h at room temperature. Nuclei were stained with 4,6-diamidino-2-phenylindole (DAPI). Fluorescence images were acquired with a Leica TCS SP2 confocal microscopy (Leica, Mannheim, Germany) using Leica Confocal Software.

### Statistical analysis

All statistical analyses were performed using SPSS 17.0 software (SPSS Inc., Chicago, IL), and graphs were generated using GraphPad Prism 5 Software (Graph Pad Software, Inc., La Jolla, California, USA). Paired t-tests, Student t-tests, and Kruskal-Wallis tests were used to analyze the mRNA and gene expression. ANOVAs were also performed to compare more than two groups. The Pearson’s correlation test was employed to evaluate the association between the expression of IGFBP5 mRNA and the disc degeneration grade of patients. P values (two-tailed) less than 0.05 were considered statistically significant.

## Acknowledgments

We thank all donors enrolled in the present study.

